# A Statistical Review of Virus Reduction in Coagulation, Flocculation, and Sedimentation Treatment Processes

**DOI:** 10.64898/2026.01.19.700160

**Authors:** Mira Chaplin, Lars Andersland, Delaney Snead, Brian M. Pecson, Charles N. Haas, Daniel Gerrity, Adam Olivieri, Tim Dinh, Avery Sanchez, James Henderson, Krista R. Wigginton

## Abstract

Coagulation, flocculation, and sedimentation (CFS) is widely applied as a combined unit process in the treatment of drinking water, wastewater, and recycled water; however, virus reduction through CFS has not been sufficiently characterized to assign pathogen log reduction value (LRV) credits. This study collected data through a systematic review that yielded over 1000 LRVs from 43 manuscripts covering 46 viruses to characterize virus reduction through CFS. The results demonstrate that CFS is effective at reducing viruses, with 68% of virus LRVs greater than 1. A mixed-effects model was used to identify potential mechanisms of virus reduction with ferric and aluminum coagulants, as well as factors associated with variability in performance. Key insights from the model show that virus reduction is: (1) improved at lower pH, similar to natural organic matter (NOM) reduction, (2) lower in secondary effluent than surface water for drinking water treatment, (3) virus-dependent, and (4) dependent on virus enumeration methods, with lower LRVs observed for molecular techniques. These findings demonstrate the potential for CFS to provide consistent and explainable virus reduction, potentially establishing a foundation for regulatory crediting in potable reuse applications. Future crediting frameworks will need to account for the factors impacting performance to accurately quantify and assign credit for virus reduction.

## 1. Introduction

The combination of coagulation, flocculation, and sedimentation (CFS) is commonly used in drinking water treatment to remove suspended solids and organics, and it also serves as an effective, and often required, pretreatment for downstream filtration and disinfection. It is ubiquitous in conventional surface water treatment plants (drinking water treatment) and is increasingly considered for advanced water treatment (AWT) trains in potable reuse applications. Beyond the typical water treatment objectives of suspended solids and organics removal, CFS has also been shown to reduce pathogens, including bacteria, viruses, and *Cryptosporidium.*^1–3^ While pathogen reduction has been demonstrated, CFS is not explicitly credited as a pathogen barrier. A primary obstacle to crediting is the fact that the extent, variability, and mechanisms of pathogen reduction through CFS remain insufficiently understood.

Although not credited on its own, CFS combined with granular media filtration does receive pathogen credit through the U.S. EPA’s Surface Water Treatment Rule.^4^ As long as filter effluent turbidity remains below 0.3 NTU in 95% of monthly measurements (40 CFR 141.173), 1 log reduction value (LRV) of virus credit is granted for direct filtration, which includes coagulation, flocculation, and filtration (without sedimentation), and 2 LRVs of virus credit are granted for conventional filtration (which includes sedimentation before filtration). This crediting framework was developed based on a dataset in which all CFS processes were immediately followed by a filter.^5^ The framework eliminated the possibility of evaluating relationships between virus reduction and water quality changes during the CFS portion of the process. This is a limitation, given that substantial virus reduction (> 1 log) has been observed from CFS alone.^6–9^

Because the drinking water regulatory framework requires the combination of CFS and filtration, it cannot be flexibly applied to novel AWT trains considered in some potable reuse applications. For example, carbon-based advanced treatment (CBAT) typically includes a combination of ozone and biofiltration, generally with biological activated carbon (BAC). The primary treatment objective for ozone-BAC is transformation and/or removal of bulk and trace organics, and these processes are not always preceded by CFS or operated as traditional filters.^10^ In these scenarios, solids removal can be achieved prior to AWT or with downstream membrane filtration processes.^11^ However, CFS is increasingly being considered for potable reuse systems to leverage its additional water quality benefits.^12^ This highlights an opportunity to seek complementary virus credits for CFS, particularly given that virus LRV targets for direct potable reuse (DPR) treatment trains are as high as 20 log in California (Cal. Code Regs Tit. 22, § 64669.45).

In effect, potable reuse has decoupled CFS and filtration, allowing for greater flexibility but also highlighting a critical regulatory crediting gap. Some potable reuse systems plan to implement the traditional drinking water framework for LRV crediting,^13,14^ but this may not be acceptable in all locations, particularly if there are differences in CFS performance in wastewater compared to surface water. Therefore, it is necessary to demonstrate the conditions under which robust virus reduction is consistently achieved, and to explore associations between virus reduction and easily measured operational and/or water quality parameters that could serve as surrogates for this potential critical control point (CCP).

Studies have documented substantial variation in virus reduction with CFS, from < 0.5 log to over 6 log.^6–9,15,16^ Scientific consensus has not yet emerged on key questions regarding the effectiveness of CFS for virus reduction, such as (1) the identification of a conservative human virus for treatment evaluation and crediting and (2) associations between virus reduction and process parameters that could serve as surrogates for assessing CFS performance. Multiple studies have demonstrated that, similarly to organic matter, increasing the coagulant dose and lowering the pH lead to greater virus reduction; however, the magnitude and range of these effects vary widely across different water matrices.^6,17,18^ Interpreting the literature is further complicated by differences in experimental design, including different viruses, coagulants and doses, the presence of coagulant aids (e.g., polymers), and reactor hydraulics. Even when studies employ similar methods, water quality differences impact CFS performance in terms of solids, organics, and viruses, making performance conclusions difficult to generalize. Because of these differences, it may not be justifiable to implement the drinking water framework for potable reuse applications. Evaluating findings on a study-by-study basis impedes our ability to identify the dominant drivers of virus reduction. A systematic review of virus reduction in CFS systems, along with an integrated statistical analysis of the resulting data, has the potential to reveal widespread patterns and develop quantitative relationships between water quality and operational parameters and virus reduction.

The decades of studies on virus reduction in CFS, combined with the lack of crediting frameworks for CFS as an independent process, motivate a systematic analysis of existing data. This study collects and models data on virus reduction in CFS systems to characterize virus reduction and understand the impact of different parameters, including operating conditions (e.g., pH, coagulant dose, coagulant type, treatment scale), virus type, and virus quantification methods (culture vs. molecular). These findings shed light on the overall ability of CFS to remove viruses independent of filtration processes. The resulting insights on the drivers of virus reduction may aid in the development of new virus reduction crediting frameworks for potable reuse applications, particularly as the industry transitions toward DPR.

## 2. Methods

### 2.1 Systematic Literature Review

We collected literature from the Engineering Village, Scopus, Water Resource Abstracts, and Web of Science databases using the Preferred Reporting Items for Systematic Reviews and Meta-Analyses (PRISMA) guidelines.^19^ In November of 2023, we searched each database using the “all fields” option with the terms “*(coagulation OR rapid mixing OR sedimentation OR flocculation OR clarification OR slow mixing) AND (water treatment) AND (virus OR bacteriophage OR coliphage) AND (removal OR inactivation OR settling)*.” Papers not in English were translated using Google Translate. Relevant references from review papers in the systematic review were added to the full-text review stage.

We screened the resulting papers using a two-stage process: abstract screening (stage 1) followed by full-text screening (stage 2). During abstract screening, one reviewer examined the title and abstract of each paper to determine if the study included the following three criteria to continue to full-text screening: (1) experiments with coagulation and flocculation treatment (excluding electrocoagulation), (2) measurements of virus or virus-like particle reduction, and (3) coagulation and flocculation conducted on impure water solution, defined as water containing material that could be coagulated. This included untreated surface and groundwater, synthetic wastewater and drinking water with added alkalinity, clay, or sand, and treated wastewater or drinking water that was not filtered through pores smaller than 0.22 μM. For cases in which it was unclear after stage 1 if the study adhered to one or more of the three inclusion criteria, we screened the full text for clarification. We note that the presence of a sedimentation step following coagulation and flocculation was not included in stage 1 criteria because this step was often indeterminable in the abstract.

Stage two included full-text screening by two reviewers to determine whether datasets in the paper satisfied additional criteria (details presented in **S1**). For this study, we define a dataset as a set of LRVs from a single paper that are presented together in a graph or table, or that share common characteristics (e.g., collected in water from the same source). In brief, we excluded papers or datasets from papers that duplicated data extracted elsewhere in the review; did not isolate the CFS virus reduction from prior or subsequent treatment; lacked a sedimentation step following coagulation and flocculation; dissolved virus flocs prior to virus quantification; had fewer than 10 infectious viruses (e.g., plaque forming units) in the untreated samples (i.e., before CFS); did not report sufficient information to calculate LRVs; did not specify the coagulant dose used in coagulation; or added viruses after coagulant addition. Because the distinction between coagulation (i.e., “rapid mix”) and flocculation was not always clear, we included papers that reported coagulation and sedimentation even if flocculation was not mentioned. For papers published after 2000 that lacked one or more required inclusion criteria, corresponding authors were contacted to request the missing information. When an article was excluded for one reason, additional exclusion criteria were not always evaluated.

### 2.2 Data Extraction and Processing

Information to calculate LRVs (e.g., initial and final concentrations, percent reduction, confidence intervals) was independently extracted by two reviewers, with LRV calculations described in **S2**. A single reviewer extracted variables from each paper (**S3**), including paper information (e.g., year, title, journal); CFS treatment variables (e.g., dose, pH, water source); CFS process descriptors (e.g., scale, coagulation speed, settling time); microbial methods used (e.g., assay details, spiking stock purification, virus information); and water quality parameters (e.g., alkalinity, dissolved organic carbon (DOC), turbidity). All extracted variables were visually reviewed by a second reviewer. Data in plots were digitized using DigitizeIt 2.5.9 software.^20^ Additional taxonomic information of the viruses used in the studies, including the virus species, genus, and family, was extracted from the International Committee for the Taxonomy of Viruses (ICTV), based on the 2024 ICTV taxonomic release.^21^

Any LRV discrepancies between the two reviewers greater than 15% (relative to the mean of the two values) were re-examined and resolved to ensure consistency (**S4**). Of the 43 LRV discrepancies initially identified between the two reviewers, 26 were identified as extraction errors and were reextracted, whereas 17 were attributed to plot resolution and both extracted values were retained. Overall, the extracted LRVs between the two reviewers were highly concordant (CCC 0.9996; **Figure S2**) and were ultimately averaged for plotting and models.

We derived several additional variables from the extracted data. When not provided in the study, coagulant doses in mg-Fe or mg-Al were calculated based on the provided dose and coagulant molecular formula (**S5**). Author clusters were defined by shared authorship, with any papers that included at least one common author assigned to the same cluster (**S6**). Water sources were grouped into seven categories, including drinking water sources, synthetic solutions, raw wastewater, secondary effluent, filtered wastewater, and spent filter backwash water (**S3**). “Drinking water sources” included untreated or partially treated drinking waters, as well as untreated surface waters. “Synthetic solutions” involved buffers with added clay, sand, or organic matter. Clay addition, sand addition, salt addition, alkalinity addition, organic addition, and organic coagulant aid addition were boolean variables defined as true when any of the relevant constituents was added, regardless of the quantity or specific formula (e.g., clinoptitolite, bentonite, and kaolin were all designated as clay). This approach was necessary because the doses of added constituents were not always provided, and some constituents were only reported in a single study. Pearson correlations were calculated between virus LRVs and water quality variables.

### 2.3 Modeling

CFS exhibits strong water-specific performance in reducing turbidity and NOM; we therefore used mixed-effects modeling to determine which factors most strongly influence virus reduction in CFS for ferric and aluminum coagulants across the literature. Treating study-level variation as random effects allows the model to explicitly acknowledge such heterogeneity, yielding cleaner estimates of the overall effects of water quality and process parameters. Mixed effects models were fit using the lme4 package, and conditional R^2^ values were calculated using the MuMin package.^22^ All model comparisons used the Akaike Information Criterion (AIC).^23^ All code used in this project is available at the following link: (https://github.com/mira-create/wigginton_coag_floc_sed_review). Model selection is described briefly below, with more details provided in **S7**.

We applied several filtering steps to remove incomparable or incomplete data prior to modeling (**Table S3**). LRVs were excluded if they were derived from censored data (i.e., instances in which final virus concentrations were below assay detection limits), if they were not measured with infectivity or molecular measurements, or if coagulants other than aluminum or ferric metals were used. Additionally, LRVs were excluded if the coagulant doses could not be converted to standard units of mg-metal/L or exceeded 100 mg-metal/L, a value selected because this is a highly improbable dose for water treatment and because > 99% of values were below this threshold (**Figure S4**). LRVs were also excluded if virus taxonomy was unclear, and if LRVs were not paired with pH values. For the remainder of the manuscript, we refer to all data extracted from the literature as the *complete data collection* and the subset selected for modelling as the *modeling data collection*. For categorical predictors with multiple levels, all levels were included or excluded together in order to avoid grouping unrelated categories. Due to substantial data reduction during filtering, some variables in the modeling data collection had fewer categories than in the complete data collection; for example, four out of the seven water source categories were absent from the modeling data collection. Degrees of freedom for categorical variables used as fixed and random effects are reported in **Table S4**.

We tested numerical and categorical variables derived from our systematic review in the models (see **S3** for details). Numerical variables related to process operations were limited to coagulant dose and pH because these parameters are widely recognized as the primary drivers of CFS performance for particulate and organic matter, and because most other numerical parameters were only present in a small subset of the extracted datasets (**Table S5; Figure S6**). To include as many samples as possible in our model, we restricted categorical variables tested to those present for more than 90% of virus LRVs.

We evaluated operational, water quality, study-level, and virus taxonomy variables as predictors of virus LRV in the modeling data collection. Most variables evaluated were operational variables, including CFS treatment scale (bench, pilot, or full), coagulation pH, coagulant, aluminum or ferric coagulant dose, and addition of organic coagulant aid. Additional water quality variables were the presence of additives, including clay, sand, salt, alkalinity, and organic matter. Additionally, we tested water source (drinking water source, secondary effluent, or synthetic water) and measurement method (molecular vs. culture). Study-level variables included year of publication, paper ID, and author cluster (groups of papers with shared authorship). Finally, virus taxonomy variables were included to differentiate viruses.

We applied several modeling steps to evaluate the impact of fixed and random effects on virus LRVs (**S7**). To determine whether a mixed model was appropriate, we fit a simple linear model using the initial set of variables described above and compared it to a mixed model with the same variables that included a random intercept for paper ID (Step 1). To identify which sources of variability should be modeled as random effects, we optimized the random effects structure (Step 2). We then tested additional random effects including coagulant dose, coagulant type, and paper cluster, where clusters represented groups of papers with shared authorship (**S6**). The model that provided the best fit, based on AIC, served as the random effects structure for the subsequent steps. To determine if virus differences were characterized by broad or specific taxonomic grouping, we compared models that incorporated different levels of taxonomy (as fixed or random effects) and selected the model with the best fit (Step 3). Finally, to ensure that all the included variables improved the fit of the model, we used stepwise backward selection to refine the fixed effects structure. The resulting explanatory model was used to assess the associations between variables in the model and virus LRV.

We developed an alternate model to identify a conservative virus for CFS using a more specific taxonomic grouping than Baltimore class. This model was based on the next-lowest AIC combination of taxonomic fixed effects and random effects identified in Step 3, and the fixed effects were further optimized as outlined in Step 4. The inclusion of specific taxonomic variables in the model allowed us to identify which viruses are most resistant to CFS, and to determine whether differences in resistance between viruses were statistically significant.

## 3. Results

### 3.1 Systematic Review Results

Of the 8,159 unique abstracts screened, 81 articles were selected for full-text review, and of these, 43 met the criteria for data extraction. During full-text review, 37 articles were excluded for the following reasons (evaluated in the following order): 4 for duplicate data with other publications; 12 where virus reduction by CFS could not be distinguished from membrane treatment or free chlorine disinfection; 15 which lacked a sedimentation step; 4 in which flocs were destroyed before LRVs were measured; 2 where untreated samples yielded fewer than 10 infectious viruses; 2 that reported only initial or final virus concentration, preventing calculation of LRV; and 3 that did not provide coagulant dose. Different datasets from the same articles were sometimes excluded for distinct reasons. For example, Lee (2017) reported LRVs for a bench-scale system without sedimentation, and for a pilot scale system that included sedimentation but where CFS LRVs were distinguishable from membrane treatment; in that case the two datasets were excluded based on distinct criteria.^24^ In several included papers, individual datasets were excluded for specific reasons detailed in **S1**.

The complete data collection included 1,625 LRVs (**Figure 1**). The 46 viruses represent 14 families in five Baltimore classes, including dsDNA, dsRNA, ssDNA, -ssRNA, and +ssRNA viruses (**Tables S10, S11**). Of the articles, 17 reported LRVs for bacteriophages or plant viruses,^18,25–40^ 13 reported LRVs for mammalian viruses, primarily those of concern for fecal-oral transmission,^7–9,15,41–49^ and 13 reported LRVs for both mammalian viruses and bacteriophages or plant viruses.^2,6,17,24,50–58^ The viruses were comprised mainly of mammalian viruses of concern for fecal-oral transmission and bacteriophage surrogates. The most common mammalian virus families were *Picornaviridae* (367 LRVs; 22.6%), including aichiviruses, coxsackieviruses, and polioviruses; *Adenoviridae* (115 LRVs; 7.1%), including adenoviruses; and *Caliciviridae* (68 LRVs; 4.2%), including caliciviruses and noroviruses. The most studied bacteriophage families were *Fiersviridae* (740 LRVs; 45.5%), containing phages f2, fr, MS2, and Qβ, and *Straboviridae* (86 LRVs; 5.3%), containing phages T2 and T3. Variable distributions in the complete data collection are characterized in **S8**.

**Figure 1.**
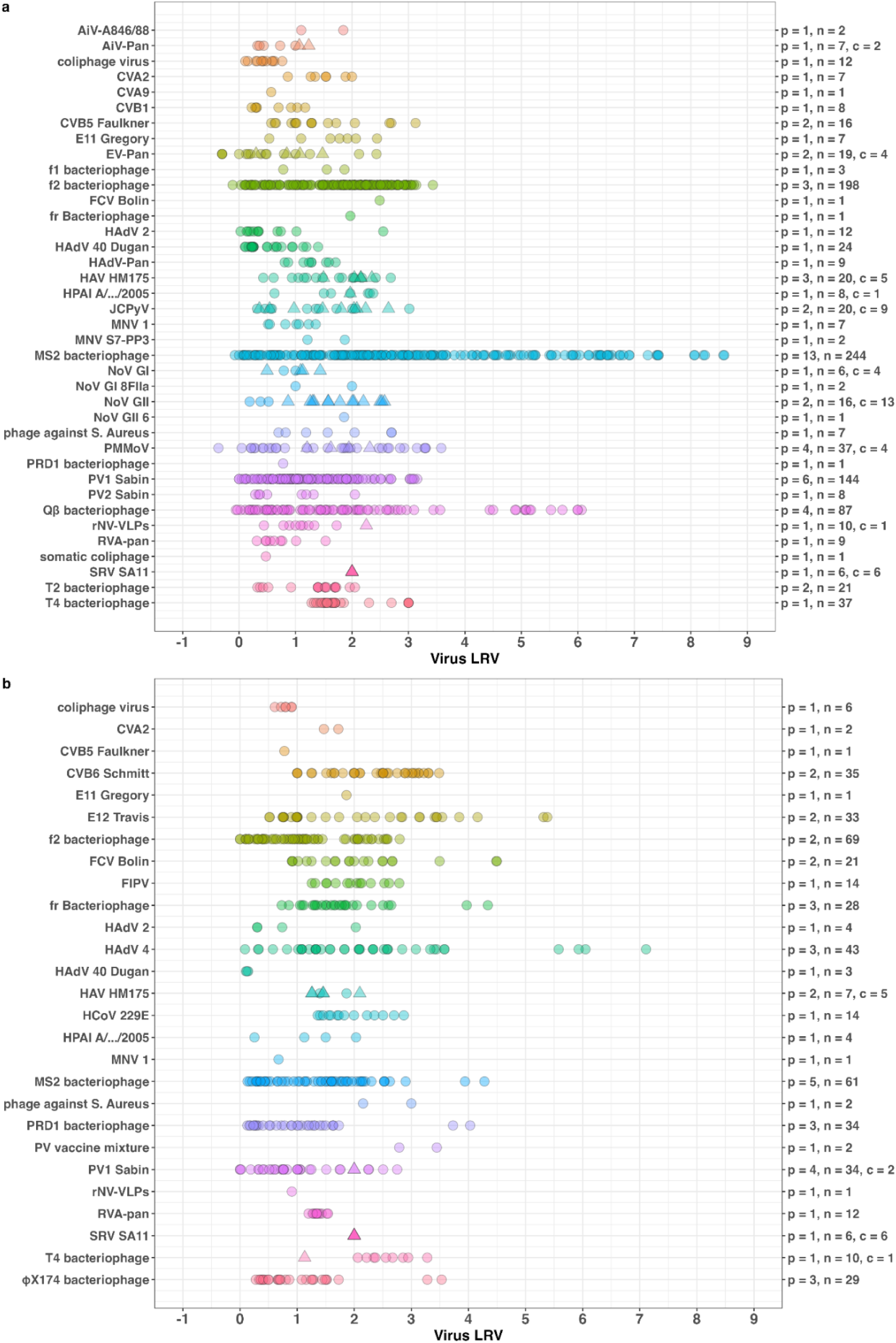
Virus LRVs collected in the systematic literature review in the complete data collection for a) aluminum and b) ferric coagulants. The secondary y-axis shows: p = number of papers, n = number of reported LRVs, and c = number of censored data points (if any). Circles = uncensored data; triangles = censored data. Point shading indicates the number of values. Virus abbreviations are defined in **Table S2**.

The data demonstrate that CFS processes alone (i.e., in the absence of a downstream filtration step) effectively reduce viruses. For example, the mean, median, and 5th percentile virus LRVs across the combined aluminum and ferric data set (excluding censored data) were 1.75, 1.52, and 0.20, respectively. Based on the empirical cumulative distribution of the data, 0.5 LRV corresponds to the 15th percentile and 1.0 LRV to the 32nd percentile (**Fig S5**). This is a significant finding, as virus reduction credit is not currently granted to CFS alone, and there is potential for substantial credit for virus reduction (e.g., > 0.5 LRV or > 1.0 LRV). These thresholds are notable because potable reuse regulatory frameworks sometimes allow crediting down to an LRV of 0.5 (18 A.A.C. 9), while others limit credited LRVs to ≥ 1.0 (Cal. Code Regs Tit. 22, § 64669.45). However, given the distribution of LRVs, with 15% below an LRV of 0.5, it is necessary to understand the water quality or operational parameters that influence LRVs and that could possibly serve as surrogates for LRV verification.

Because it is impractical to directly measure the reduction of viruses through CFS, a crediting framework would ideally be based on an easily measured surrogate parameter that is highly correlated with virus reduction. However, in the complete data collection, single-parameter correlations between LRV and water quality or operational variables (pH, alkalinity, conductivity, DOC, TOC, turbidity, UV_254_ absorbance, coagulant dose, and pH) were uniformly weak (all Pearson coefficients < 0.31; **Table S5**). These results are unsurprising given that single-parameter correlations with water quality or operational variables do not account for differences in coagulant type, virus family, and source water characteristics. Weak correlations indicate that virus reduction in CFS relies on complex mechanisms that cannot be explained by one single overarching parameter. However, when the data were limited to specific viruses, coagulants, or water sources to evaluate the suitability of single-parameter models under more limited conditions, the remaining data sets were too sparse to support reliable correlation analyses. These limitations motivated our use of multivariable models to capture the combined and interacting effects of coagulant dose, water quality, and virus characteristics on virus reduction in CFS.

### 3.2 Model Selection Results

The goal of our modeling effort was to identify the water quality and operational parameters consistently associated with greater virus reduction, thus providing insights that might inform CFS design, operation, and regulatory crediting. To build the mixed-effects model, we first narrowed from 1,625 LRVs in the complete data collection to 1,050 LRVs in the modeling data collection by removing LRVs that did not include comparable coagulant doses and pH, among other factors (**Table S3**). We evaluated multiple models with different combinations of variables and used AIC to identify the model that best balanced data fit and parameter complexity for explaining virus reduction in CFS.

The final explanatory model (AIC = 2,672, conditional R^2^ = 0.77; Eq. 1) had 34 fixed-effect parameters, including both continuous and categorical predictors. The continuous predictors were coagulation pH and log_10_(coagulant dose). The remaining predictors were categorical and included Baltimore virus class, coagulant type, use of coagulant aid, measurement method (molecular vs. culture), treatment scale (bench, pilot, or full), and water quality (secondary effluent, surface water, or synthetic water). This model also included several random effects to account for systematic differences among the studies, coagulants, and viruses. These consisted of random intercepts for each paper, random intercepts for each virus within each paper, and random intercepts and slopes for log_10_(coagulant dose) within each paper. We note that additional variables that may influence virus reduction, such as alkalinity, DOC, TOC, turbidity, and UV_254_ absorbance, could not be included due to the sparsity of reported data across the reviewed studies. The inability to include these variables likely contributes to the need for the complex random effects structure.

The structure of the final explanatory model is shown in equation 1:

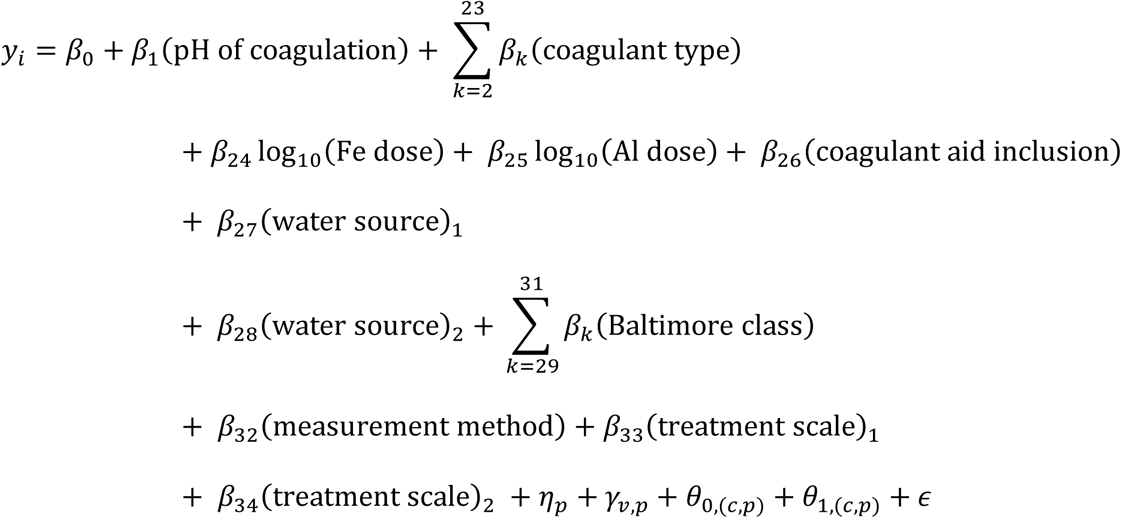

where *y_i_* is the LRV for data point *i*, β values are model coefficients, ε is the unobserved residual error, and *p(i)*, *v(i)*, and *c(i)* are the paper ID, virus abbreviation, and coagulant for data point *i*, respectively, with the dependence on *i* suppressed to enhance readability. Mixed-effects models represent random effects as drawn from underlying probability distributions with parameters estimated from the data. The distributions and parameters of the model random effect are shown in equations 2-4:

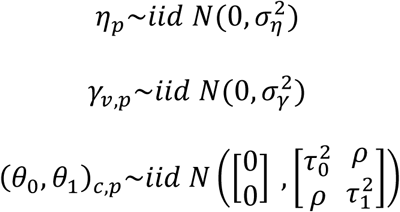

where 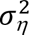 is the variance of the paper random effects, 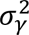 is the variance of the virus x paper random effects, 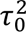 is the variance of the random intercepts for coagulant times paper, 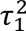 is the variance of the random slopes for coagulant dose by paper and coagulant, and ρ is the covariance between the intercept and slope within a paper and coagulant. Distributions of variables in the model are presented in **Tables 1 and2**.

**Table 1.** Distributions of continuous variables present in the final explanatory model, separated by aluminum coagulants and ferric coagulants. Coagulant doses were converted into standard units of mg-metal/L.

|  | Aluminum |  |  |  | Ferric |  |  |  |
| --- | --- | --- | --- | --- | --- | --- | --- | --- |
| Variable | mean | std | min | max | mean | std | min | max |
| coagulant dose*<br>(mg-metal/L) | 2.3 | 2.0 | 0.3 | 15.8 | 14.8 | 1.9 | 2.2 | 58.9 |
| log10(coagulant dose<br>(mm-metal/L)) | 0.4 | 0.3 | -0.5 | 1.2 | 1.2 | 0.3 | 0.4 | 1.8 |
| coagulation pH | 6.9 | 0.8 | 5.2 | 8.4 | 6.6 | 0.8 | 5.1 | 9.2 |

\*included in the model as log10(coagulant dose)

**Table 2.**
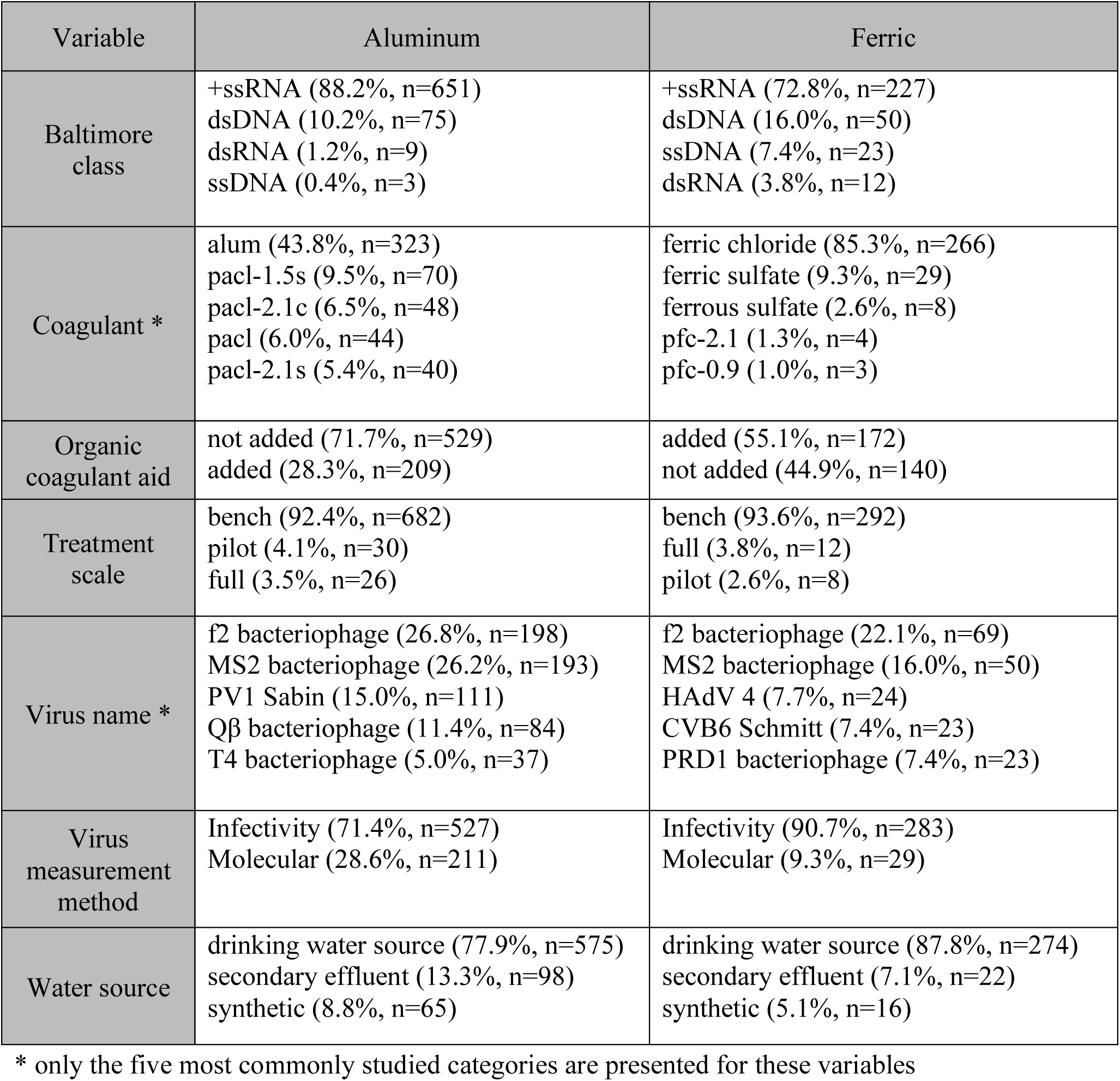
Distributions of categorical variables in the final explanatory model. The percentage and n values signify the proportion and number of samples in the specified category, respectively.

The final explanatory model included only variables that improved model performance (i.e., lowered AIC). Some categorical variables included levels that were not individually significant (**Figure 2**). For example, synthetic water was not statistically different from the drinking water reference for virus reduction, but secondary effluent was statistically different. As illustrated in Figure 2, higher LRVs were associated with lower coagulation pH, higher coagulant doses, surface waters compared to secondary effluent, +ssRNA viruses compared to ssDNA and dsDNA viruses, culture-based virus measurements compared to molecular-based virus measurements, pilot-scale and full-scale studies compared to bench-scale systems, and the addition of organic coagulant aids. In the following sections, we examine each of these variables, as well as the random effects. We discuss their distribution in the modeling data collection, define their estimated effects, explore potential mechanistic explanations grounded in previous literature, and consider how our findings may inform system design, operation, and optimization. We also highlight remaining uncertainties and data gaps that limit our understanding and identify areas of future research.

**Figure 2.**
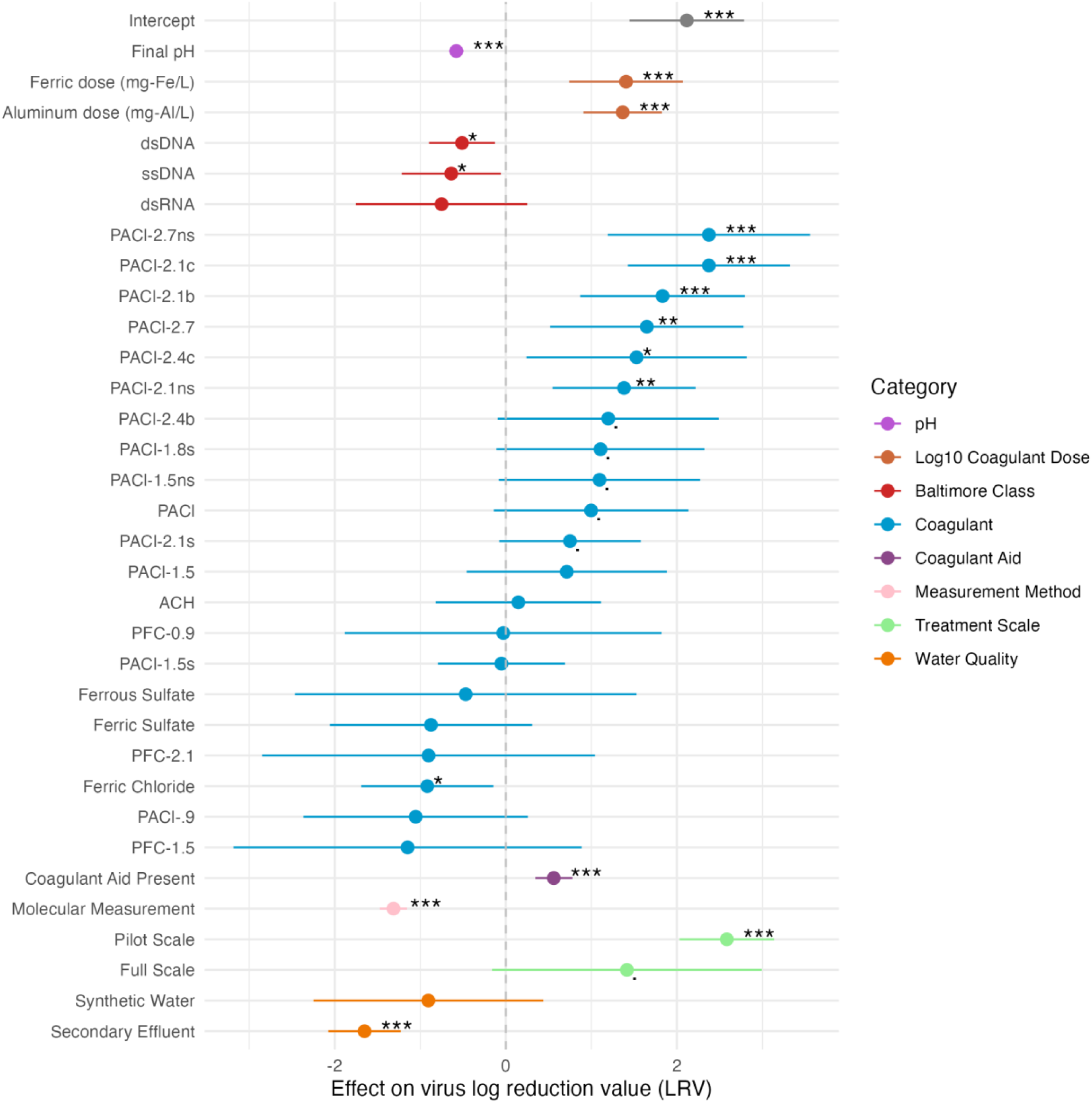
Coefficients for categorical and numerical variables in the final explanatory model. Asterisks indicate statistical significance (“.”: < 0.1; * : < 0.05; **: < 0.01; ***: < 0.001). Points illustrate the impact of the variable on model LRV estimates relative to a reference category or value of the variable; the addition of these coefficient values to the model intercept provides the estimated LRV. The categorical variable reference categories are: +ssRNA for the Baltimore class variable, aluminum sulfate for the coagulant type variable, coagulant aid absent for the coagulant aid variable, infectivity for the measurement variable, bench scale for the treatment scale variable, and drinking water source for the water quality variable. For the numerical variables, log_10_(coagulant dose) was referenced to 0 (coagulant dose of 1 mg-metal/L), and pH was referenced to pH 5.

### 3.2.1 Random Effects

Including paper-specific random effects allows the model to capture distinct chemical interactions between the unique water matrices used in each paper and viruses, coagulants, and coagulant doses without biasing the fixed-effect estimates or overfitting limited data. The final explanatory model accounted for 77% of the variance in virus reduction (conditional R^2^ = 0.77), which is a strong fit given the diversity of the modeling data collection. The inclusion of random effects substantially improved fit; the model using only fixed effects explained 37% of the variance (marginal R^2^ = 0.37). Models including the same fixed effects but with only a subset of random effects captured much of this additional variance. A model with paper-level random intercepts explained 72% of the variance (conditional R^2^ = 0.72), while a model that also included a random intercept for coagulant explained 75% of the variance (conditional R^2^ = 0.75). The addition of a random slope for dose produced a model that explained 76% of the variance (conditional R^2^ = 0.76). The random intercept for virus within a paper yielded only a modest increase in conditional R^2^ (0.76 to 0.77) but improved model AIC, indicating residual within-study heterogeneity among viruses that is not captured by fixed effects alone.

The adjusted intraclass correlation coefficient (aICC) is a metric to evaluate the proportion of the total variance, excluding the fixed effects, explained by each random intercept.^59^ Because the final explanatory model includes a random slope, the aICC values are dependent on the units of the log_10_(coagulant dose), and therefore cannot be considered as an overall diagnostic of the model. Nonetheless, the relative proportions of the aICC values of the random intercepts are constant and can be used to compare the contributions of the random intercepts.

The random intercept for paper ID explained a large portion of the model variance unexplained by fixed effects relative to the other random intercepts (aICC = 0.24), likely reflecting differences in water composition between treatment plants, inherent measurement variability, or measurement error. The differences in water quality could be attributable to unmodeled parameters that are correlated within treatment plants but were reported sparsely in the modeling data collection, such as temperature or NOM concentration, as well as additional parameters not reported in any studies. Smaller proportions of the variance were explained by the random intercepts for coagulant within a paper (aICC = 0.09) and virus within a paper (aICC = 0.06). The performance of CFS for turbidity and NOM reduction are known to be highly specific to individual treatment plants, and these modest paper-specific differences in LRV for individual viruses and coagulants demonstrate that virus reduction is similarly variable. While the aICC of the random slope for coagulant dose within coagulant and paper is not comparable to the other aICC values, its improvement of the model AIC demonstrates that the relationship between coagulant dose and virus LRV varies between treatment plants, similar to the relationship between coagulant dose and turbidity or NOM reduction. Taken together, these random effects substantially improved model fit and captured key sources of between-study variability, allowing more robust estimation of the fixed effects.

### 3.2.2 pH

pH is frequently measured and calibrated in CFS, especially to optimize NOM reduction in enhanced coagulation. It is therefore important to evaluate the effect of pH on virus reduction.

In our final explanatory model, lower pH was strongly associated with greater virus reduction (β = -0.58, p < 0.001), indicating that LRVs are estimated to decrease by 0.58 units for every 1-unit increase in pH starting at the reference pH of 5. These model results reflect the findings of individual studies. Our modeling data collection included 66 data subsets (containing 538 LRVs) in which pH varied within a single paper while other variables were held constant (**S10**). Lower pH was associated with greater virus reduction in 60 of these 66 subsets. Coagulant addition usually lowers pH, but the association between lower pH and greater virus reduction remained even when the coagulant dose was held constant (14 subsets) or was not significantly correlated with LRV (26 subsets), suggesting that pH is associated with virus reduction independent of coagulant dose. Because NOM reduction is also enhanced at lower pH values,^60^ DOC and UV_254_ deserve attention as potential surrogates for virus reduction.

These subsets with pH variation within a single paper provide an opportunity to investigate mechanisms that potentially link virus reduction to pH. Mayer (2015) posited that virus reduction could increase at pH values far from the virus isoelectric point (IEP), a virus-specific parameter referring to the pH value at which a virus is neutrally charged.^61^ To explore this, we compared the LRV-pH slopes of subsets reported for viruses with distinct IEPs. For this analysis, 26 subsets with significant LRV-dose correlations (p < 0.1) were first removed because coagulant addition both lowers pH and leads to greater virus reduction, confounding pH-specific effects (**S10**). Linear regressions of LRV versus pH were fit for the remaining 40 subsets. Among these, 37 were conducted entirely at pH values above the virus IEP. Of the 37 subsets, 33 exhibited negative slopes, indicating higher LRVs at lower pH even though lower pH was closer to the virus IEP. The remaining three subsets (for poliovirus, coxsackievirus, and echovirus, **Table 2**) included pH values that were mostly, but not exclusively, above the virus IEP. These subsets showed similarly negative LRV-pH relationships rather than low LRVs near the virus IEP.

These results differ from the proposed hypothesis and suggest that IEP alone does not consistently predict how virus reduction varies with pH. Although negative slopes were common, their magnitudes differed significantly, even for the same virus. This does not preclude a role for IEP; instead, its influence may depend on interaction with other virus-specific properties, NOM properties, and aggregation state. To determine the mechanisms behind the influence of pH on different viruses, researchers should consistently report water quality parameters, particularly NOM concentration and characteristics. Predictive models incorporating such parameters with virus characteristics could improve our understanding of reduction mechanisms.

**Table 2:** Viruses studied, number of subsets per virus, and reported versus selected isoelectric points (IEPs) for the 40 subsets where pH varied within a single paper while other variables were held constant. Subsets with significant LRV-dose correlations (p < 0.1) were removed. “Reported IEP ranges” lists virus IEPs reported in reviews and primary literature,^62–65^ while “Analysis IEP” indicates the representative value selected for this study (**S10**).

| **Virus** | **Number of subsets** | **Reported IEP ranges** | **IEP used in analysis** |
| --- | --- | --- | --- |
| human adenovirus 4 | 1 | 2.6 | 2.6 |
| MS2 bacteriophage | 17 | 2.2, 3.1-3.9 | 3.5 |
| pepper mild mottle virus | 1 | 3.7-3.8 | 3.7 |
| PRD1 bacteriophage | 2 | 3.8-4.2 | 3.9 |
| f2 bacteriophage | 6 | 4 | 4 |
| Qβ bacteriophage | 10 | 1.9, 2.7, 5.3 | 5.3 |
| echovirus (echovirus 12 in this study) | 1 | 5-6.4 | 5.7 |
| coxsackievirus (coxsackievirus B6 Schmitt in this study) | 1 | 4.75-4.8, 6.1, 6.75 | 5.8 |
| poliovirus 1 Sabin | 1 | 4.5, 6.5 | 6.2 |

### 3.2.3 Water Source

CFS is most commonly used for drinking water treatment but is also increasingly of interest for treating secondary effluent in potable reuse trains.^12^ Understanding whether virus reduction is consistent across water matrices is essential for extrapolating findings across treatment applications (e.g., drinking water vs. wastewater) and for developing future regulatory frameworks for virus reduction in potable reuse systems. In our modeling data collection, 849 (80.9%) LRVs were measured in untreated drinking water sources, 120 (11.4%) were measured in secondary effluent, and 81 (7.7%) were measured in synthetic water. In the final explanatory model, secondary effluent showed lower LRVs than drinking water sources (β = –1.65, p < 0.001), indicating that CFS is less effective at removing viruses in wastewater effluent than drinking water sources under otherwise equivalent conditions. The model findings align with the findings from the single paper in the data collection that studied CFS in both drinking water sources and wastewater. That study reported substantially higher LRVs in untreated surface water than in secondary effluent when using the same coagulant doses for coagulation of f2 bacteriophage, but did not provide any hypothesis for these discrepancies.^38^

A potential reason for the higher LRVs observed in drinking water sources compared to wastewater is the generally greater NOM content and different NOM characteristics in secondary effluent compared to drinking water influent.^66^ Higher NOM concentrations have been linked to decreased virus reduction through coagulation treatment processes.^16,42,67^ Shamsollahi (2019) reported a strong inverse correlation between initial TOC concentration and rotavirus reduction over a year of CFS operation. Similarly, Tanneru et. al (2012) reported 2-4 log lower MS2 reduction in synthetic water with added NOM during electrocoagulation-microfiltration. A proposed explanation for these observations is that NOM interferes with coagulation by competing with viruses for coagulant binding sites, shielding virus surfaces, or stabilizing viruses in suspension. As described above, IEP does not appear to be a primary driver of virus reduction, and NOM and viruses seem to demonstrate similar pH dependence, suggesting competition between viruses and NOM viruses and NOM might exist. Variations in NOM composition could further explain reduced virus LRVs in secondary effluent compared to surface water. NOM in secondary effluent is typically more hydrophilic than NOM in surface water,^68^ and coagulation removes hydrophilic NOM less effectively than hydrophobic NOM.^69^ The more persistent NOM in secondary effluent rather than drinking water sources may limit virus reduction through virus competition with NOM for active sites.

Although virus reduction by CFS is generally lower in secondary effluent than in surface water, substantial virus reduction (LRVs up to 6.5) has been demonstrated for bacteriophages in secondary effluent under optimized treatment conditions.^24,30,33,38,39^ However, the two available studies of human virus reduction in secondary effluent show variable results, including some LRVs below 0.5.^24,49^ Because of the substantial differences between virus reduction in drinking water and secondary effluent, direct adoption of the drinking water regulatory framework or use of drinking water data to create regulations for secondary effluent risks overestimating virus LRVs. Additional research will be necessary to determine whether CFS achieves consistent reduction of human viruses in secondary effluent and to identify the process parameters to ensure consistent performance.

### 3.2.4 Viruses

Understanding whether different viruses are removed differently in CFS is essential for the design of robust treatment systems and for the selection of appropriate surrogates for viruses of concern. Although 22 papers in the complete data collection compared the reduction of different viruses across identical conditions, they frequently demonstrated inconsistent rank orders of virus susceptibility. For example, human adenovirus was more resistant to CFS than other human viruses in some studies,^15,41^ but was removed similarly or more effectively than enteroviruses, caliciviruses, and rotaviruses in other studies.^43,50^ Because many other factors differ among the studies, the cause of the observed inconsistencies cannot be easily identified. Mixed-effects modeling provided an opportunity to identify trends between viruses across the entire literature that were not evident when comparing individual studies to one another.

Virus taxonomy poses challenges for mixed-effects modeling because viruses can be classified at several taxonomic levels, each of which can be expressed in models as a fixed or random effect. We tested multiple combinations of taxonomic variables as fixed effects, random intercepts, or random intercepts nested within paper ID (**Table S7**). The final explanatory model suggests study-specific variability in the behavior of individual viruses (represented by virus name as a random intercept crossed with paper ID), while demonstrating that overall, cross-study patterns in virus reduction are most consistently reflected at the level of Baltimore class (represented as a fixed effect). With respect to trends between virus groupings, the dsDNA (β = - 0.51, p = 0.006) and ssDNA (β = -0.64, p = 0.024) viruses showed lower LRVs than the reference group of +ssRNA viruses. Although viruses in the same Baltimore class share little beyond their genome structure, virus size is a possible explanation for this pattern. The dsDNA viruses in the modeling data collection (adenoviruses, tectiviruses, straboviruses, and polyomaviruses) are generally larger than the +ssRNA viruses in the data collection, which could influence their reduction mechanisms during coagulation.^61,64^ However, the ssDNA viruses (microviruses and inoviruses) are similarly sized to the +ssRNA viruses in the data collection, so size is not an explanation for their resistance to CFS. Overall, these results suggest there is likely no single mechanism associated with the relative resistance of the different Baltimore classes.

To further explore differences between viruses, we evaluated a supplementary model using virus family as a fixed effect and the interaction between virus name and paper ID as a random effect (**Figure S8**). All other virus families had lower estimated LRVs than the reference family Picornaviridae, although the confidence intervals of the coefficients are large (**Figure 4**). Only two families were statistically different from Picornaviridae: Microviridae (β = -0.67, p = 0.038) and Tectiviridae (β = -0.75, p = 0.020). In the modeling data collection, Microviridae included only the common surrogate bacteriophage φX174, and Tectiviridae included only the bacteriophage PRD1. These findings, in addition to data excluded from the model due to lack of coagulant dose information,^15^ suggest that bacteriophage φX174 is conservative relative to important waterborne human viruses.

**Fig 4.**
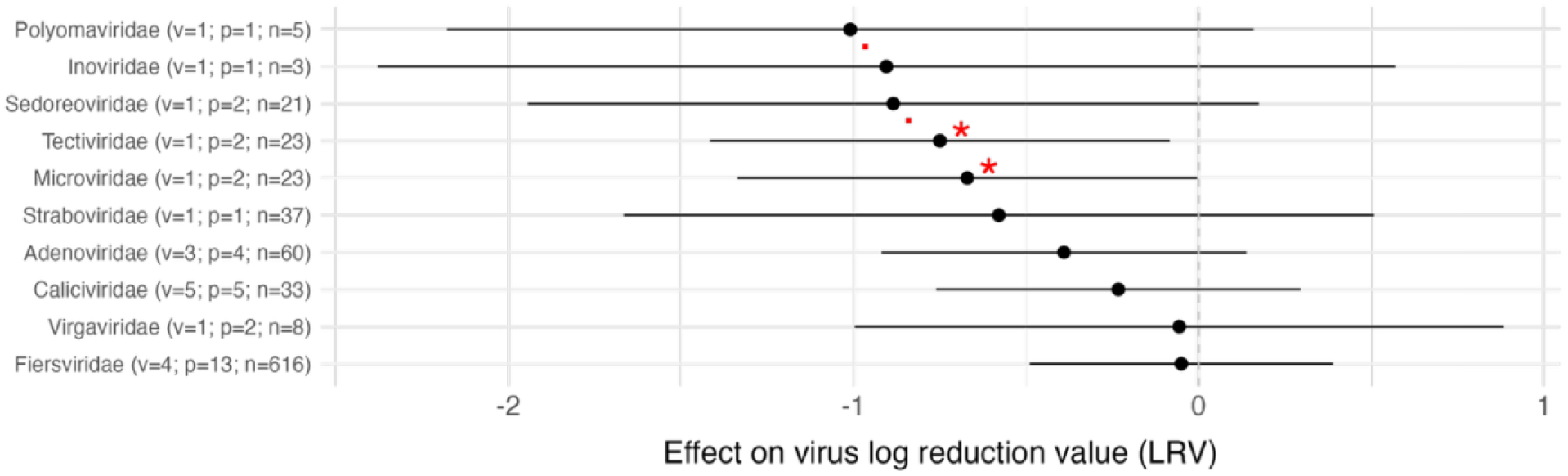
Coefficients for virus family in the supplementary taxonomic model. (“.”: p < 0.1; * : p < 0.05). The reference virus family is Picornaviridae. v refers to the number of viruses in the family, p refers to the number of papers that studied the virus, and n refers to the number of reported LRVs.

### 3.2.5 Measurement Method

Virus reduction in water treatment is typically measured using either culture-based assays (810 points; 77.1% of the modeling data collection) or molecular assays (240 points; 22.9% of the modeling data collection), each of which has distinct advantages and limitations. For example, culture-based assays may not distinguish a single infectious virus particle from a virus aggregate and therefore might overestimate virus reduction when viruses aggregate during CFS treatment. Likewise, molecular assays do not indicate whether the detected viruses are infectious, potentially leading to underestimation of virus reduction when viruses are inactivated through CFS treatment. Understanding how these methods compare is critical for interpreting reduction data, identifying biases, and developing reliable crediting frameworks.

Our results suggest that culture-based assays show higher virus LRVs through CFS treatment. In the final explanatory model, virus reduction measured using molecular methods was substantially lower than reduction measured using culture-based assays (β = 1.31, p < 0.001). This may reflect either virus inactivation or aggregation of infectious viruses. To explore the extent and sources of this difference in LRV based on assay type, we examined the six studies in the complete data collection that directly compared molecular and infectivity measurements collected from experiments done with the same viruses, coagulant, dose, water matrix, and pH (**Figure 5**).^18,24,36,46,47,52^ These studies demonstrated moderate correlation between the two methods (Pearson coefficient = 0.56), with exceptions for polyaluminum chlorides used with MS2 and Qβ

**Fig 5.**
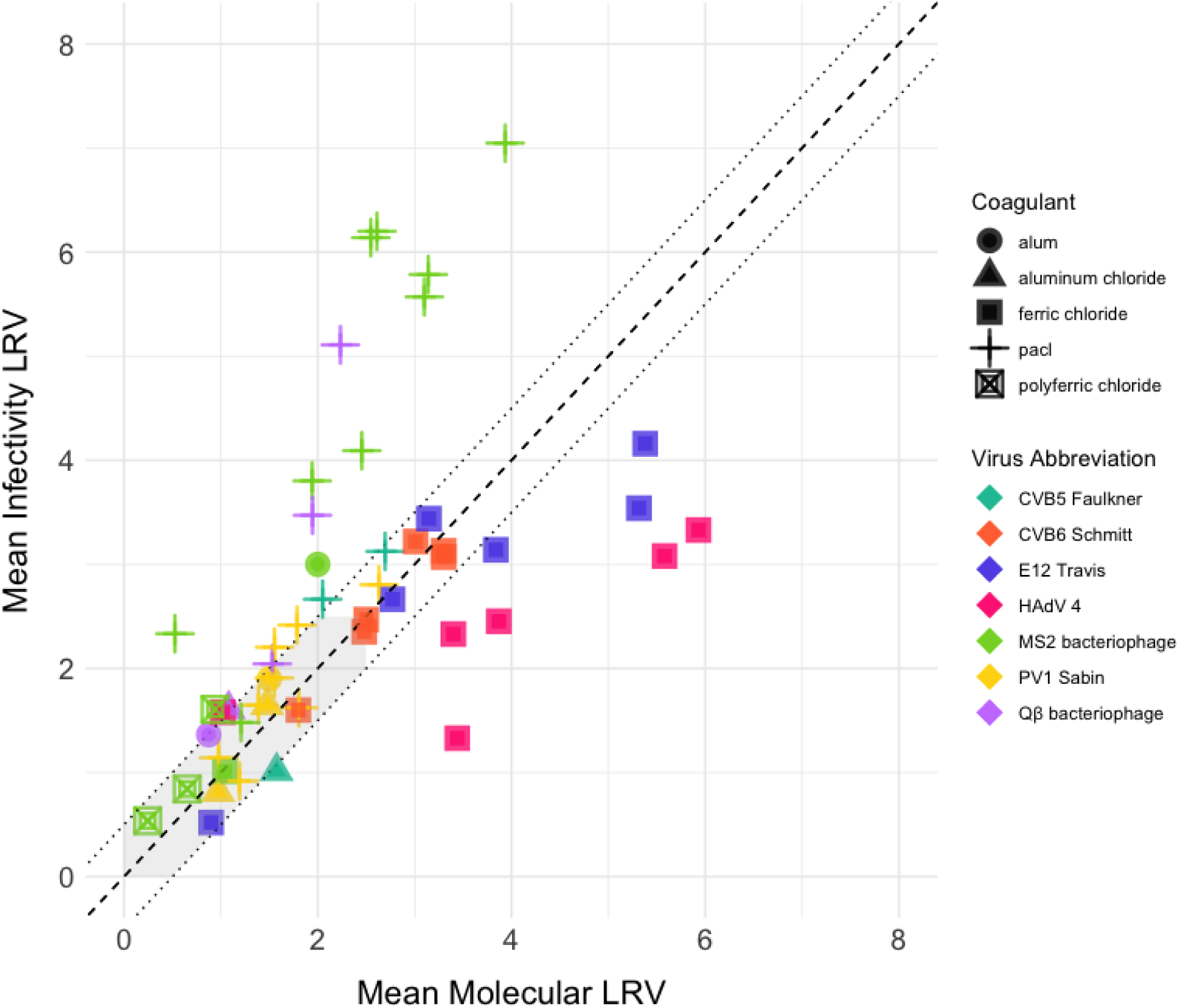
Mean culture-based LRVs plotted against mean molecular-based LRVs for studies in the complete data collection where the same virus was studied at the same conditions (coagulant, dose, water, and pH). None of these studies used a coagulant aid. Replicates are averaged, including replicates at similar, but not identical, pH (defined as pH values where the author targeted a specific pH, but measured slightly different final pH values [<0.2 pH unit difference]).

and ferric chloride used with HAdV2 and E12 Travis. The original studies attributed the PACl discrepancies to virus inactivation with PACl,^36,47^ and the ferric chloride discrepancy to the underestimation of virus reduction using the cytopathic effect assays for human adenovirus.^46^

These results suggest that no single measurement approach is universally appropriate for evaluating virus reduction in CFS. The magnitude of the difference between infectivity and molecular measurements likely depends on the interactions between the virus, coagulant, and water source. Further research should investigate this difference for additional human viruses and explore the extent to which the difference is due to inactivation or aggregation. Because virus concentration and quantification methods can be biased by changes in the water matrix,^58^ we recommend that future studies report methodological recoveries in both untreated and treated samples.

### 3.2.6 Treatment Scale

Bench-scale studies are commonly used to evaluate virus reduction during coagulation processes, especially when pilot- or full-scale testing is logistically or financially impractical. For turbidity and NOM reduction, bench-scale results are often used to inform design decisions, validate full-scale performance, and support regulatory crediting, but it is unclear whether bench-scale data accurately reflect virus reduction in larger systems. Our final explanatory model results demonstrate that pilot-scale systems generally result in higher LRVs (β = 2.58, p <0.001) than bench-scale systems. Full-scale systems also exhibit higher LRVs than bench-scale systems (β = 1.42, p = 0.065).

Results from the individual studies generally reinforce the statistical models’ findings that virus reduction is greater in full- and pilot-scale systems than in bench-scale systems. Rao (1988) and Abbaszadegan (2007) both described greater reduction in full-scale than bench-scale without offering potential explanations for the discrepancy.^17,45^ Nearly all of the bench-scale studies in the modeling data collection use spiked viruses, whereas full- and pilot-scale studies more frequently study indigenous viruses present in the source water. Indigenous viruses in surface water or wastewater may belong to different clades than spiked viruses and/or may enter CFS already associated with the wastewater matrix (e.g., particles) and therefore settle differently. Differences in source water quality, particularly for full-scale studies relying on secondary effluent or low-turbidity water, may also contribute to lower observed LRVs. Taken together these findings suggest that bench-scale studies are valuable for exploring treatment mechanisms, identifying conservative LRVs and relative trends, but full-scale testing provides the most accurate estimate of actual virus reduction through CFS operations.

## 4. Conclusions

This study compiles and synthesizes decades of data across a range of viruses, water matrices, coagulants, and operational conditions to characterize virus fate during CFS. By combining a systematic literature review with a statistical model fit using over 1000 observations, we provide one of the most comprehensive quantitative evaluations of virus reduction during CFS to date. Our findings offer practical insights for treatment system design, validation, and crediting by identifying broad trends that are often obscured in individual studies. This model could be used to support a virus crediting framework in CFS for both drinking water and potable reuse applications.

Model insights are useful for selecting surrogates, assigning virus reduction credit, and guiding future research on the mechanisms responsible for virus reduction in treatment systems. CFS is not currently granted virus LRV credits separately from filtration; however this study shows that significant virus reduction is achieved in CFS. The model could be used to identify the conditions, coagulant doses, and pH values needed to consistently achieve certain LRV thresholds, such as the minimum LRVs that must be demonstrated to justify credit based on existing potable reuse regulatory frameworks. The model demonstrates that (1) virus reduction is lower in secondary effluent than in surface water, potentially justifying different frameworks for potable reuse vs. conventional drinking water; (2) virus reduction is substantially greater at low pH; (3) φX174 and PRD1 are the viruses most consistently resistant to CFS, highlighting their potential role as conservative surrogates in future studies; and (4) virus reduction is lower when measured using molecular methods.

Several variables were not included in our model. Temperature, flocculation time, mixing intensity, sedimentation time, initial pH, and alkalinity and other parameters all have either a demonstrated or plausible impact on CFS. These parameters were excluded because they were reported with low precision or were missing from a significant portion of the data collection. Including these parameters in future models could improve the fit and further clarify the mechanistic drivers of virus reduction. Our modeling data collection is significantly smaller than the initial data collection (1050 vs 1625 rows) and also contains less variation in key variables. Our exclusion of 63 censored data points from the model could make our model predictions overly conservative, particularly for molecular measurements of human viruses. Despite these limitations, the model captures consistent, interpretable trends and provides a foundation for future predictive tools and crediting frameworks.

## Acknowledgements

This work was supported by Environmental Protection Agency STAR #84025601. M. C. was supported by a National Science Foundation Graduate Research Fellowship (award no. #2241144).

## Declaration of generative AI

During the preparation of this work the authors used ChatGPT in order to prepare figures and code. After using this tool, the authors reviewed and edited the content as needed and take full responsibility for the content of the published article.

