## Supplemental Material 1 - 10 for "A Statistical Review of Virus Reduction in Coagulation, Flocculation, and Sedimentation Treatment Processes"

List of Supplementary Exhibits

**S1**. Methods for systematic review and data extraction

**S2**. Details on calculation of log reduction values

**S3.**Variables tested in the models

**S4.** Comparison of LRVs between reviewers

**S5.** Coagulant dose conversions

**S6.** Cluster analysis

**S7.** Model selection

**S8.** Variable distributions in the complete data collection

**S9.** Model details

**S10.** pH analysis

**S11**. Complete data collection from systematic review

### **S1.** Methods for systematic review and data extraction

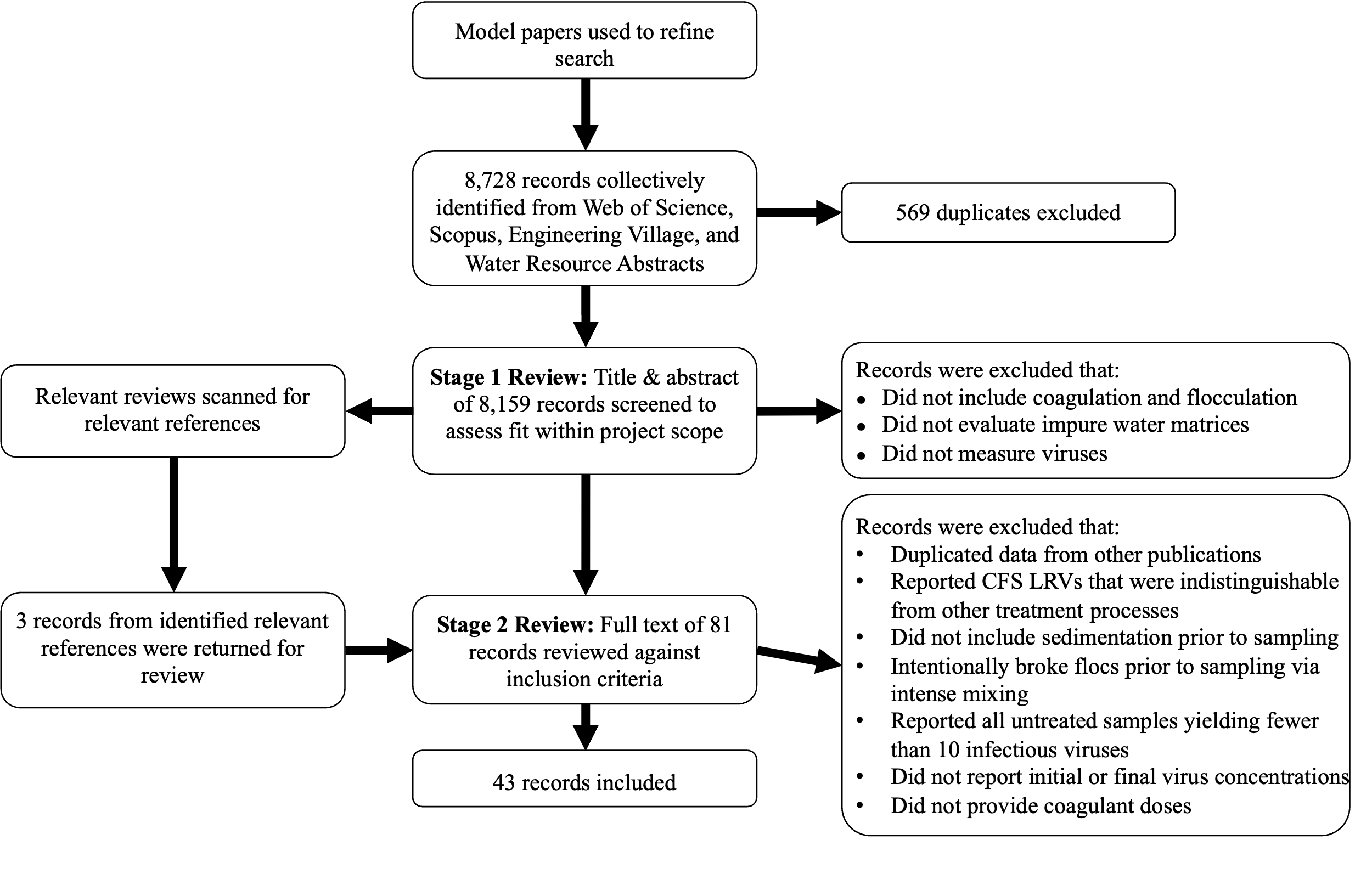

#### **Figure S1**. Flowchart for systematic review. The two-stage review resulted in 43 articles.

We evaluated the papers using a two-stage screening process. During the abstract screening, one reviewer examined the title and abstract of each paper to determine whether it reported: (1) coagulation and flocculation, (excluding electrocoagulation), (2) measurement of virus reduction, and (3) impure water solution. Impure water was defined as water containing material that could be coagulated, such as natural waters; synthetic wastewater and drinking water with added coagulation material; and treated wastewater or drinking water that had not been filtered through pores smaller than 0.22 𝜇M. Ultrapure solutions, distilled water, and phosphate buffers without added alkalinity were classified as pure solutions.

In this second stage of review, samples were rejected if:

1. They duplicated data present in other publications by the same authors. In this case, only the most comprehensive and detailed version was retained, usually the paper with the earliest publication date. (4 articles excluded)
2. Reported CFS LRVs were indistinguishable from virus reduction due to disinfection, filtration, or other treatment processes. Datasets reporting process-specific CFS LRVs in treatment trains with subsequent or prior treatment processes were included, with the exception of datasets with chlorine disinfection prior to CFS. In these cases, a chlorine residual likely contributes to subsequent virus reduction in CFS. (12 articles excluded)
3. Sedimentation was not clearly described, did not occur, or if sampling took place during flocculation rather than after settling. (15 articles excluded)
4. Flocs were intentionally broken up prior to sampling via intense mixing. This practice deviates from the typical CFS operation and could skew virus reduction data. (4 articles excluded)
5. Untreated samples yielded fewer than 10 viral colonies, as this was considered insufficient for quantifying reduction following CFS. (2 articles excluded)
6. Only the initial or final virus concentration was reported, so the LRV could not be calculated. (2 articles excluded)
7. No coagulant dose was provided. (3 articles excluded)

Some datasets from included articles were excluded for additional reasons:

8. Viruses added to water after coagulant addition, as this is a highly nonphysical scenario;
9. When PCR inhibitor removal treatment was used, the same treatment had to be applied to the samples before and after CFS;
10. Reported a coagulant dose of zero;
11. Virus reduction was measured after centrifugal separation of floc mixture.

### **S2.** Details on calculation of log reduction values

Parameters and LRVs were extracted from text, tables, and figures. When both tables and figures were provided, LRVs were preferentially extracted from tables due to higher precision.

When LRVs were directly reported, those values were extracted. Where LRVs were not directly provided, LRVs were calculated based on initial and final virus concentration or percent reduction as summarized in Table S1.

#### **Table S1**. Calculation methods for virus log reduction value (LRV) when not directly reported.

| **Calculation Method** | **Equation** | **Paper references** |
| --- | --- | --- |
| Initial (N0) and final (N) Concentration | $log_{10}(\frac{N_{0}}{N})$ | ^1–8^ |
| Percent reduction (%) | -$log_{10}(\frac{\%}{100})$ | ^9–17^ |

### **S3.**Variables tested in the models

Below is a description of all variables tested in the models. Reference levels used in the model are provided for categorical variables tested as fixed effects. Additional variables extracted from the paper but not tested in the model are described in the uploaded dataset.

*Paper ID*: ID number assigned during review *(categorial, no reference)*

*Cluster***:** Papers were grouped into clusters connected by common authors. (see S6) *(categorial, no reference)*

*Virus Taxonomy (virus name, family, species, Baltimore class)*: The Baltimore class of a virus is general knowledge. Virus species and family were determined based on the International Committee for the Taxonomy of Viruses 2024 taxonomic release.^18^ Viruses were assigned a unique name based on the most granular possible differentiation, which was frequently below the species level. Virus names and abbreviations correspond to standard naming and conventions and are provided in Table S2.

Viruses measured using molecular methods were frequently characterized by the PCR assay used to measure them. Different PCR assays were combined when they were used to characterize viruses in the same taxonomic group, with the same level of specificity. For example, different PCR assays to measure pan norovirus GII were given the same virus name, but an assay measuring only norovirus GII (6) was given a distinct name. *(all categorical, reference virus name: MS2 bacteriophage, reference species: Emesvirus zinderi, reference family: Fiersviridae/Picornaviridae (depending on model), reference Baltimore class: +ssRNA)*

#### **Table S2.** Virus abbreviations used throughout the paper

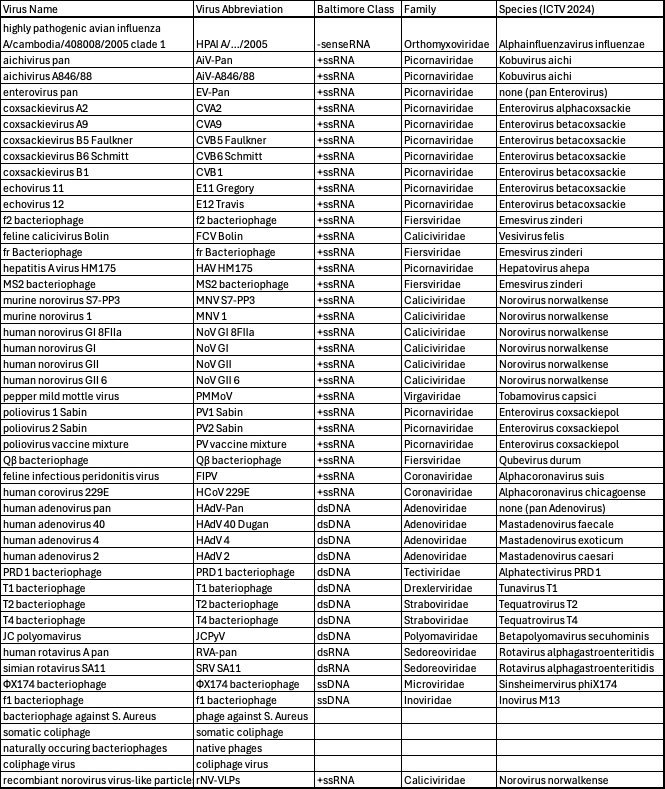

*Treatment Scale*: The treatment scale refers to the size and implementation of the study, divided into bench, pilot, or full. Full-scale indicated operational treatment systems. Pilot-scale studies replicated operational conditions at reduced scale. Bench-scale trials were defined as controlled lab tests, often conducted as jar tests. *(categorical, reference = bench scale)*

*Water Type*: CFS source waters were grouped into 7 categories. Detailed source water descriptors (e.g., source river or lake) were extracted when available. For categories, “drinking water source” included untreated or minimally treated drinking water sources, even if blended (e.g., surface + groundwater). We combined all surface waters, as it was often unclear whether drinking water treatment plant influent referred to surface water, groundwater, or a variable blend of the two. “Treated drinking water” refers to tap or fully treated sources. Synthetic source refers to lab solutions starting with ultrapure water or buffer and with added clay, sand, or organic matter. Additional categories are raw wastewater, secondary effluent, filtered wastewater, and spent filter backwash water. *(categorical, reference = drinking water source)*

*Year of Publication:* determined from the paper, and referenced to 1958, the earliest year of a study in the modeling data collection. *(continuous, reference = 1958)*

*Water Quality Amendment (clay addition, sand addition, salt addition, alkalinity addition, organic addition, and organic coagulant aid addition):* presence/absence variables defined as true when any of the relevant constituent was added, regardless of the quantity or precise formula (e.g. clinoptitolite, bentonite, and kaolin were all designated as clay). *(categorical, reference = no addition)*

*Coagulant type* : Coagulant names reported in each study were recorded. Coagulants were differentiated as long as they had different molecular formulas (e.g., different molecular formulas of polyaluminum chloride were set as different variables). Coagulants with the same composition of metals and salts were deemed the same coagulant, regardless of waters of hydration (e.g. FeCl3 and FeCl3*6H20 were both designated as ferric chloride). *(categorical, reference = aluminum sulfate)*

*Virus measurement method* : Technique used for virus quantification. This is divided into two main categories: infectivity (plaque assay, TCID50, mouse assay) and molecular (qPCR, ddPCR). *(categorical, reference = infectivity)*

*pH after treatment*: During coagulation, the pH drops rapidly with the addition of coagulant and then stabilizes during sedimentation and flocculation. In some cases, pH was recorded post-treatment without indication of adjustment. In others, pH was manually adjusted prior to the experiment to achieve a specific target final value through the addition of strong acids or bases.^13,19–21^ When the paper described a “coagulation pH” or stated that coagulation happened at a certain pH, this was assumed to be the final pH. Sometimes, pH was adjusted to achieve a target initial value; when the final pH value was not reported, this initial value was not deemed sufficient to provide information on the coagulation pH. *(continuous, reference = pH 5)*

*Log_10_ coagulant dose:* converted to mg-metal/L of Fe or Al, as described in S5 *(continuous, reference = 0 (coagulant dose of log_10_[1 mg-metal/L])*

### **S4.** Comparison of LRVs between reviewers

Following data extraction, all LRV differences between reviewers exceeding a 15% threshold were flagged for manual review. The reviewers evaluated whether each discrepancy reflected an extraction error or arose from low absolute values combined with limited plot resolution (e.g., the difference between 0.5 LRV and 0.6 LRV is >15%, but may not be further resolvable if the LRVs are small when compared to the axis of a plot). If extraction errors were identified, reviewers re-extracted the data.

Following these steps, Lin’s concordance correlation coefficient was calculated to assess agreement across all data points. Using >0.99 as an acceptable target, the resulting coefficient of 0.999601 confirms a high degree of consistency between reviewers and supports the overall reliability of the extracted data collection, as visualized in Figure S2.

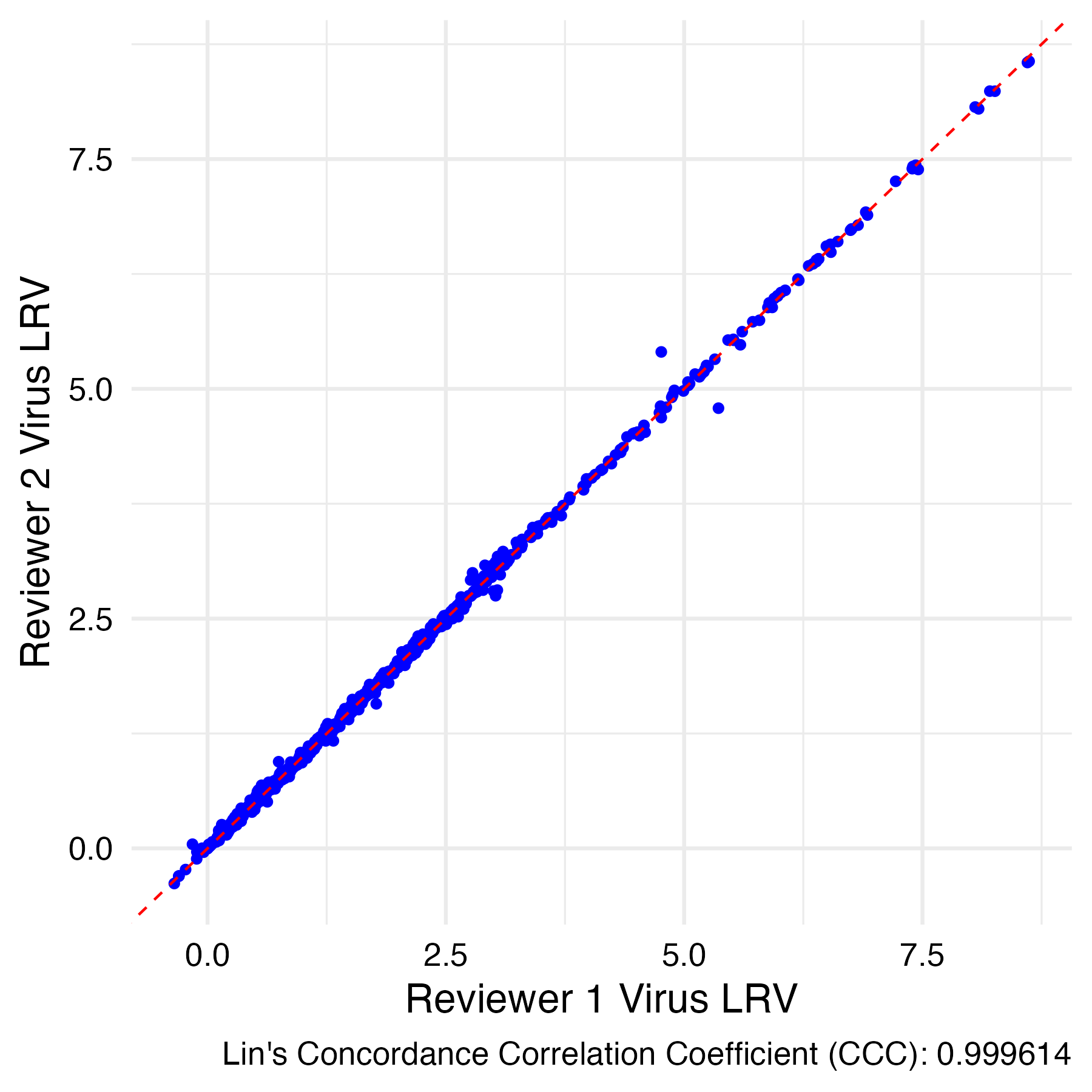

#### **Figure S2**. Agreement between reviewers in data extraction. Lin’s concordance correlation coefficient >0.99 indicates strong agreement between reviewers.

### **S5.** Coagulant dose conversions

To ensure consistency across studies, coagulant doses were standardized using molecular weight conversions for each ferric or aluminum coagulant with a provided molecular formula (e.g., FeCl₃, Fe₂(SO₄)₃, Al₂(SO₄)₃·18H₂O). For example, a dose of 20 mg Fe₂(SO₄)₃ was converted to mg of elemental Fe by (1) dividing by the molar mass of Fe₂(SO₄)₃, (2) multiplying by the stoichiometric ratio of Fe atoms, and (3) multiplying by the molar mass of Fe, as shown:

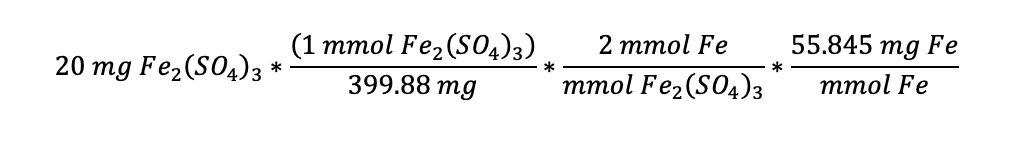

Hydrated species were handled explicitly by including the molar mass of water in the compound weight. When coagulant doses were reported in micromolar metal units (e.g., µM-Fe), conversions to mg-metal/L were also performed directly via atomic weights.

When the molecular formula was not available, dose in mg-metal/L was not calculated and the samples were excluded from modeling. Although dose was a required inclusion criterion, the coagulant molecular formula was not available for at least some coagulants in 9 papers. All of the excluded samples were for aluminum-based coagulants, including aluminum sulfate,^17,22–25^ polyaluminum chloride (PACl),^5,23,26^ or aluminum chloride.^27^ All assumptions, logic, and unit mappings are documented in the dose standardization functions within the data cleaning script.

### **S6.** Cluster analysis

To assess potential author-related biases, a cluster analysis was performed by mapping shared authorship across studies. Each node in the resulting graph in Figure S3 represents a paper, and edges indicate shared authorship between papers. Most data points appear as relatively isolated nodes, suggesting minimal bias overlap. Significant clusters were defined as those with three or more interconnected papers; three such clusters were identified.

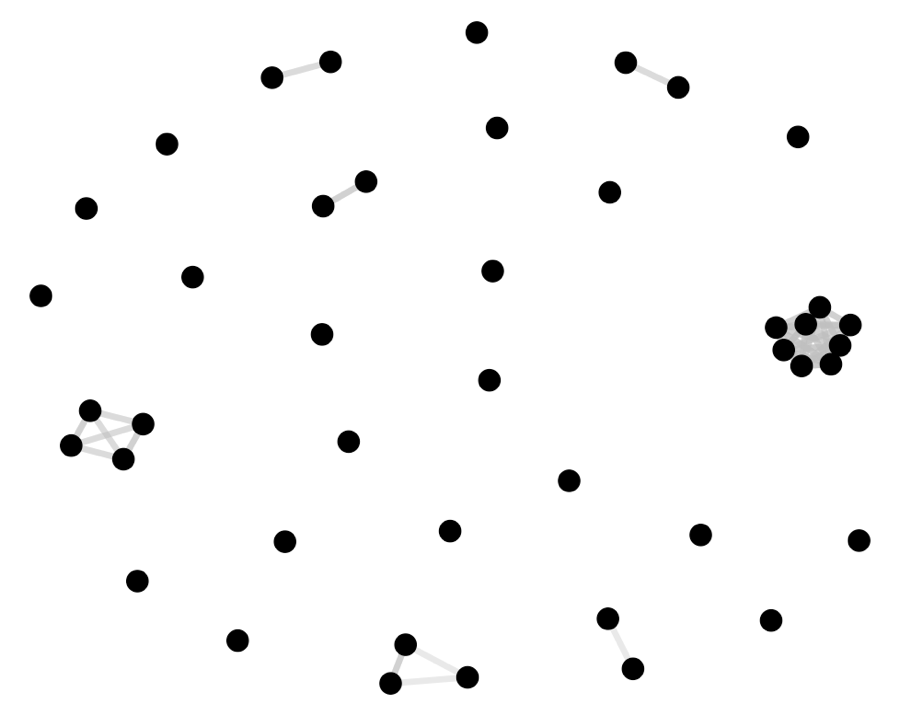

#### **Figure S3**. Cluster diagram for shared authorship between papers included in the systematic review. Nodes represent publications and lines represent shared authors.

### **S7.** Model selection

### **Table S3.** Data filtered during preparation for modeling

| **Operation** | **Number of LRVs in data collection** | **Number of papers in data collection** | **Coagulant distribution** | **Number of LRVs dropped** |
| --- | --- | --- | --- | --- |
| Complete data collection | 1625 | 43 | Aluminum: 1021  Ferric: 477 |  |
| Remove censored rows | 1562 | 43 | Aluminum: 972  Ferric: 463 | 63 |
| Remove rows where virus measurement method is neither infectivity or molecular | 1552 | 43 | Aluminum: 963  Ferric: 462 | 10 |
| Remove coagulants that are not ferric or aluminum | 1425 | 39 | Aluminum: 963  Ferric: 462 | 127 |
| Removed coagulants that could not be converted | 1298 | 31 | Aluminum: 836  Ferric: 462 | 127 |
| Remove rows with coagulant dose > 100 mg-metal/L | 1296 | 31 | Aluminum: 836  Ferric: 460 | 2 |
| Remove rows with undefined taxonomy (i.e., coliphage virus, phage against S. Aureus) | 1281 | 30 | Aluminum: 829  Ferric: 452 | 15 |
| Drop rows with no pH | 1050 | 20 | Aluminum: 738  Ferric: 312 | 231 |

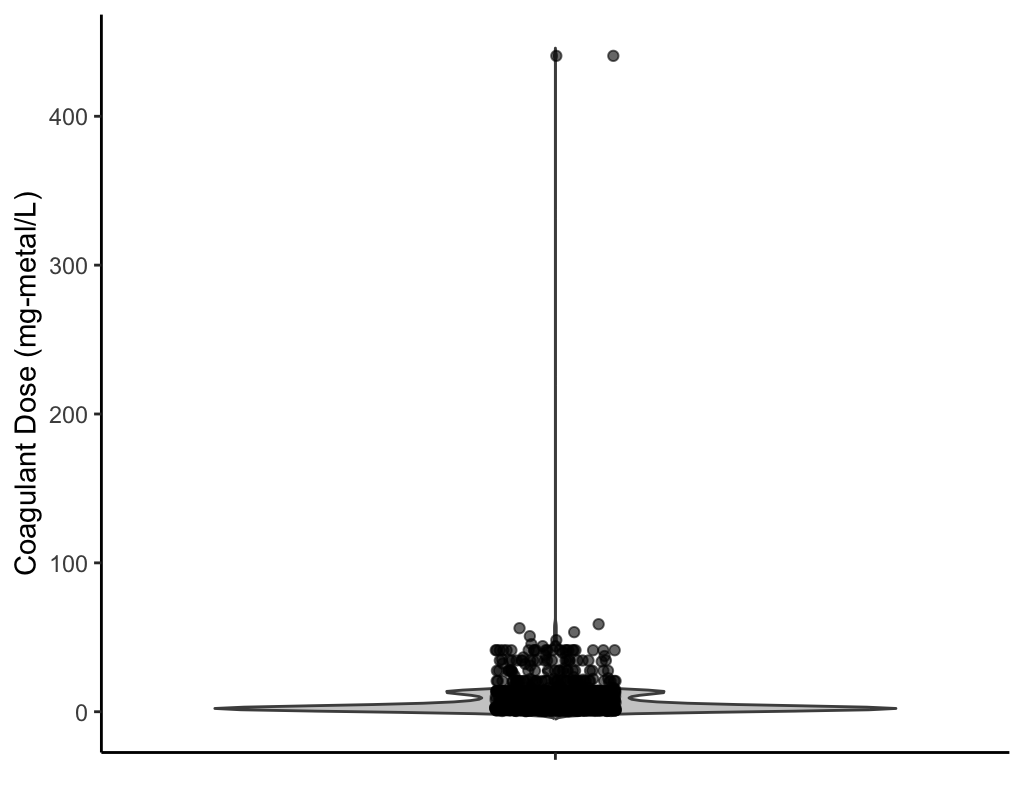

#### **Figure S4**. Distribution of coagulant doses (mg-metal/L) in data collection prior to removal of rows where coagulant dose > 100 mg-metal/L

#### **Table S4.** Degrees of freedom of categorical variables in the modeling data collection

| Variable | Degrees_of_Freedom (unique values - 1) |
| --- | --- |
| Record ID | 19 |
| Cluster | 9 |
| Virus Name | 27 |
| Species | 19 |
| Family | 10 |
| Baltimore Class | 3 |
| Treatment Scale | 2 |
| Water Type | 2 |
| Publication Year | 13 |
| Clay addition | 1 |
| Sand addition | 1 |
| Alkalinity addition | 1 |
| Organic adding | 1 |
| Organic coagulant aid addition | 1 |
| Coagulant type | 21 |
| Virus measurement method | 1 |

**Table S5**. Pearson correlations of virus LRV with initial, final, and percent reduction values for water quality parameters, along with the number of contributing studies.

| **Parameter** | **Pearson correlation coefficient** | **R^2^ of correlation** | **Number of papers reporting parameter** |
| --- | --- | --- | --- |
| Alkalinity Initial (mg-CaCO3/L) | -0.18 | 0.03 | 18 |
| Alkalinity Final (mg-CaCO3/L) | -0.18 | 0.03 | 2 |
| Alkalinity % Reduction | 0.31 | 0.09 | 2 |
| Conductivity Initial  (units variable) | -0.18 | 0.03 | 6 |
| Conductivity Final  (units variable) | -0.20 | 0.04 | 4 |
| Conductivity % Reduction | 0.03 | 0.001 | 4 |
| DOC Initial (mg/L) | -0.10 | 0.01 | 13 |
| DOC Final (mg/L) | -0.03 | 0.001 | 2 |
| DOC % Reduction | 0.28 | 0.08 | 2 |
| TOC Initial (mg/L) | -0.30 | 0.09 | 4 |
| TOC Final (mg/L) | -0.11 | 0.01 | 1 |
| TOC % Reduction | -0.20 | 0.04 | 1 |
| Turbidity Initial  (units variable) | -0.14 | 0.02 | 25 |
| Turbidity Final  (units variable) | -0.20 | 0.04 | 8 |
| Turbidity % Reduction | 0.29 | 0.09 | 8 |
| UV254 Initial (cm^-1^) | -0.18 | 0.03 | 5 |
| UV254 Final (cm^-1^) | -.014 | 0.02 | 2 |
| UV254 % Reduction | 0.09 | 0.09 | 2 |
| Initial pH | -0.09 | 0.01 | 23 |
| Final pH | -0.08 | 0.01 | 24 |
| Coagulant Dose | -0.10 | 0.01 | 43 |

The steps of the modeling approach are as follows:

1. **Linear vs mixed model with paper ID comparison:** A simple linear model was fit to the modeling data collection using virus name, treatment scale, water type, year, coagulant, clay addition, sand addition, alkalinity addition, organic addition, presence of organic coagulant aid, final pH, measurement method, aluminum coagulant dose, and ferric coagulant dose. This model was compared to a mixed model fit on the same data collection with a random intercept added for the paper ID.
2. **Random effect structure optimization:** The mixed model from Step 1 was compared to two alternative models: one with a random intercept for author cluster (groups of papers with the same authors), and one with both paper and cluster random intercepts together. Building on the optimal model, we tested random effects structures including: (1) a random intercept for each paper ID-coagulant combination, (2) a random slope for coagulant dose within each of these combinations, and (3) both the random slope and intercept together.
3. **Taxonomic variable structure optimization:** We compared the optimal model from step 2 to models with virus name, family, and species as fixed effects, as well as models with each taxonomic category as random intercepts and as random intercepts crossed with paper ID. We tested every combination of the above variables, while allowing each model to have no more than one taxonomic fixed effect and one taxonomic random effect. We tested taxonomic variables as random effects because many viruses had very small amounts of data, and it seemed therefore plausible that more evidence would be required to determine the difference from the mean.
4. **Stepwise backward selection of fixed effects:** We applied a stepwise procedure to refine the potential predictors (excluding virus name). Starting with the optimized random effects model of Step 3, we removed the fixed effect from the model whose exclusion resulted in the largest AIC decrease, or in an AIC increase of no greater than 0.5. This process was repeated iteratively until removing additional fixed effects no longer improved the AIC.

**Table S6.** Model improvement during steps 1 and 2 of model selection. Green rows indicate the rows with the lowest AIC for each model selection step

## .

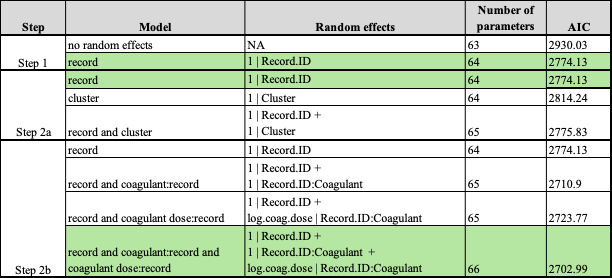

#### **Table S7.** Model improvement during step 3 of model selection. The green row is the model with the lowest AIC used for the main model. The blue row is the model used for the supplementary taxonomic model.

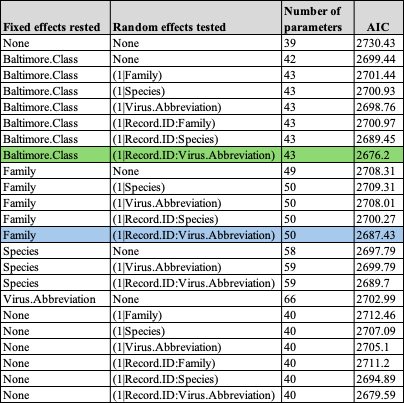

#### **Table S8.** Model improvement during step 4 of model selection

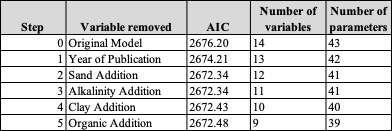

#### **Table S9.** Model improvement during step 4 of taxonomic model selection

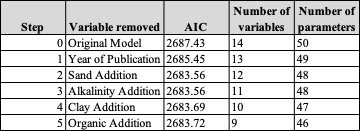

### **S8.** Variable distributions in the complete data collection

The extracted LRVs spanned a range of experimental scales, measurement methods, coagulants, and water sources (**Tables S9, S10**). Most of the LRVs (1,438; 88.5%) were derived from bench-scale systems, while 95 (5.8%) and 92 (5.7%) were obtained from full-scale and pilot-scale systems, respectively. Infectivity measurements accounted for 1,246 LRVs in the data collection (76.7%), molecular measurements accounted for 368 LRVs (22.6%), and an additional 11 LRVs were obtained with enzyme-linked immunosorbent assays to measure virus-like particles. Virus LRVs were measured across a range of water sources. A total of 1,120 LRVs (68.9%) were measured in drinking water sources, 174 LRVs (10.7%) in secondary wastewater effluent, and 198 (12.2%) in synthetic buffers. The remaining LRVs in the data collection were measured in other natural and partially treated waters: treated drinking water (86 LRVs; 5.3%), raw wastewater (32 LRVs; 2.0%), filtered wastewater (14 LRVS; 0.9%), and filter backwash water (1 LRV, 0.1%). Over 90% of LRVs were reported from treatments using aluminum or ferric coagulants. Specifically, 1,021 (62.8%) of the virus LRVs were collected from aluminum coagulants, including aluminum sulfate, aluminum chloride, and polyaluminum chlorides, and 477 (29.4%) were collected from ferric coagulants, including ferric chloride, polyferric chlorides, ferric sulfate, and ferrous sulfate. An additional 127 (7.8%) were collected with miscellaneous coagulants, including formulated coagulant aids, chitosan acetate, and lime. Our modeling focused on aluminum and ferric coagulants; however, virus reduction (median log reduction > 1 LRV) was also reported with chitosan acetate, lime, and Primafloc C-7 in individual papers. Of all the 1,624 virus LRVs, 63 (3.88%) were censored.

#### **Table S10.** Distribution of taxonomic variables in complete data collection

| **Category** | **# of LRVs** | **Percentage of LRVs** |
| --- | --- | --- |
| **Baltimore Class** | | |
| +ssRNA | 1240 | 76.3 |
| dsDNA | 280 | 17.2 |
| dsRNA | 33 | 2.0 |
| ssDNA | 32 | 2.0 |
| -ssRNA | 12 | 0.7 |
| MISSING | 28 | 1.7 |
| **Family** | | |
| Fiersviridae | 740 | 45.5 |
| Picornaviridae | 367 | 22.6 |
| Adenoviridae | 115 | 7.1 |
| Straboviridae | 86 | 5.3 |
| Caliciviridae | 68 | 4.2 |
| Virgaviridae | 37 | 2.3 |
| Tectiviridae | 35 | 2.2 |
| Sedoreoviridae | 33 | 2.0 |
| Microviridae | 29 | 1.8 |
| Coronaviridae | 28 | 1.7 |
| Drexlerviridae | 24 | 1.5 |
| Polyomaviridae | 20 | 1.2 |
| Orthomyxoviridae | 12 | 0.7 |
| Inoviridae | 3 | 0.2 |
| MISSING | 28 | 1.7 |
| **Species** | | |
| Emesvirus zinderi | 653 | 40.2 |
| Enterovirus coxsackiepol | 197 | 12.1 |
| Enterovirus betacoxsackie | 106 | 6.5 |
| Qubevirus durum | 87 | 5.4 |
| Tequatrovirus T4 | 61 | 3.8 |
| Norovirus norwalkense | 46 | 2.8 |
| Mastadenovirus exoticum | 43 | 2.6 |
| Tobamovirus capsici | 37 | 2.3 |
| Mastadenovirus caesari | 36 | 2.2 |
| Alphatectivirus PRD1 | 35 | 2.2 |
| Rotavirus alphagastroenteritidis | 33 | 2.0 |
| Sinsheimervirus phiX174 | 29 | 1.8 |
| Hepatovirus ahepa | 27 | 1.7 |
| Mastadenovirus faecale | 27 | 1.7 |
| Tequatrovirus T2 | 25 | 1.5 |
| Tunavirus T1 | 24 | 1.5 |
| Vesivirus felis | 22 | 1.4 |
| Betapolyomavirus secuhominis | 20 | 1.2 |
| none (pan Enterovirus) | 19 | 1.2 |
| Alphacoronavirus chicagoense | 14 | 0.9 |
| Alphacoronavirus suis | 14 | 0.9 |
| Alphainfluenzavirus influenzae | 12 | 0.7 |
| Enterovirus alphacoxsackie | 9 | 0.6 |
| Kobuvirus aichi | 9 | 0.6 |
| none (pan Adenovirus) | 9 | 0.6 |
| Inovirus M13 | 3 | 0.2 |
| MISSING | 28 | 1.7 |

#### **Table S11.** Distribution of viruses in complete data collection

| **Category** | **# of LRVs** | **Percentage of LRVs** |
| --- | --- | --- |
| **Virus Name** | | |
| MS2 bacteriophage | 330 | 20.3 |
| f2 bacteriophage | 294 | 18.1 |
| PV1 Sabin | 183 | 11.3 |
| Qβ bacteriophage | 87 | 5.4 |
| T4 bacteriophage | 61 | 3.8 |
| HAdV 4 | 43 | 2.6 |
| PMMoV | 37 | 2.3 |
| HAdV 2 | 36 | 2.2 |
| CVB6 Schmitt | 35 | 2.2 |
| PRD1 bacteriophage | 35 | 2.2 |
| E12 Travis | 33 | 2.0 |
| fr Bacteriophage | 29 | 1.8 |
| ΦX174 bacteriophage | 29 | 1.8 |
| HAdV 40 Dugan | 27 | 1.7 |
| HAV HM175 | 27 | 1.7 |
| T2 bacteriophage | 25 | 1.5 |
| T1 bateriophage | 24 | 1.5 |
| FCV Bolin | 22 | 1.4 |
| RVA-pan | 21 | 1.3 |
| JCPyV | 20 | 1.2 |
| EV-Pan | 19 | 1.2 |
| coliphage virus | 18 | 1.1 |
| CVB5 Faulkner | 17 | 1.0 |
| NoV GII | 16 | 1.0 |
| FIPV | 14 | 0.9 |
| HCoV 229E | 14 | 0.9 |
| CVB1 | 12 | 0.7 |
| HPAI A/.../2005 | 12 | 0.7 |
| PV2 Sabin | 12 | 0.7 |
| SRV SA11 | 12 | 0.7 |
| rNV-VLPs | 11 | 0.7 |
| CVA2 | 9 | 0.6 |
| HAdV-Pan | 9 | 0.6 |
| phage against S. Aureus | 9 | 0.6 |
| E11 Gregory | 8 | 0.5 |
| MNV 1 | 8 | 0.5 |
| AiV-Pan | 7 | 0.4 |
| NoV GI | 6 | 0.4 |
| f1 bacteriophage | 3 | 0.2 |
| AiV-A846/88 | 2 | 0.1 |
| MNV S7-PP3 | 2 | 0.1 |
| NoV GI 8FIIa | 2 | 0.1 |
| PV vaccine mixture | 2 | 0.1 |
| CVA9 | 1 | 0.1 |
| NoV GII 6 | 1 | 0.1 |
| somatic coliphage | 1 | 0.1 |

#### **Table S12.** Distribution of treatment variables in complete data collection

| **Category** | **# of LRVs** | **Percentage of LRVs** |
| --- | --- | --- |
| **Treatment Scale** | | |
| bench | 1438 | 88.5 |
| full | 95 | 5.8 |
| pilot | 92 | 5.7 |
| **Measurement Category** | | |
| Infectivity | 1246 | 76.7 |
| Molecular | 368 | 22.6 |
| Misc | 11 | 0.7 |
| **Water Type** | | |
| drinking water source | 1120 | 68.9 |
| synthetic | 198 | 12.2 |
| secondary effluent | 174 | 10.7 |
| treated drinking water | 86 | 5.3 |
| raw wastewater | 32 | 2.0 |
| filtered wastewater | 14 | 0.9 |
| spent filter backwash water | 1 | 0.1 |

#### **Table S13.** Distribution of coagulants in complete data collection

| **Category** | **# of LRVs** | **Percentage of LRVs** | **Coagulant type** |
| --- | --- | --- | --- |
| **Coagulant** | | | |
| alum | 507 | 31.2 | aluminum |
| ferric chloride | 405 | 24.9 | ferric |
| pacl | 106 | 6.5 | aluminum |
| pacl-1.5s | 78 | 4.8 | aluminum |
| ferric sulfate | 51 | 3.1 | ferric |
| pacl-2.1c | 48 | 3.0 | aluminum |
| pacl-2.1s | 42 | 2.6 | aluminum |
| pacl-2.1b | 34 | 2.1 | aluminum |
| aluminum(iii) chloride hexahydrate | 32 | 2.0 | aluminum |
| pacl-2.1ns | 25 | 1.5 | aluminum |
| pacl-2.7 | 24 | 1.5 | aluminum |
| cat floc (cationic) | 19 | 1.2 | misc |
| pacl-1.5ns | 17 | 1.0 | aluminum |
| pacl-2.7ns | 17 | 1.0 | aluminum |
| water aluminum chloride (wac) | 17 | 1.0 | aluminum |
| clinoptilolite | 16 | 1.0 | misc |
| pacl-1.5 | 16 | 1.0 | aluminum |
| pacl-2.4c | 16 | 1.0 | aluminum |
| pacl-1.8s | 15 | 0.9 | aluminum |
| pacl-2.4b | 15 | 0.9 | aluminum |
| primafloc c-7 | 15 | 0.9 | misc |
| ba-2 | 14 | 0.9 | misc |
| catfloc | 9 | 0.6 | misc |
| pei-k | 9 | 0.6 | misc |
| drewfloc # 21 (cationic) | 8 | 0.5 | misc |
| ferrous sulfate | 8 | 0.5 | ferric |
| chitosan acetate | 6 | 0.4 | misc |
| lime | 5 | 0.3 | misc |
| strychnos potatorus solution | 5 | 0.3 | misc |
| acquapol 893/11 | 4 | 0.2 | misc |
| acquapol cl 18 | 4 | 0.2 | misc |
| acquapol of 18 | 4 | 0.2 | misc |
| acquapol t832 | 4 | 0.2 | misc |
| acquapol ww | 4 | 0.2 | misc |
| ferrous alum | 4 | 0.2 | both |
| ferrous chloride | 4 | 0.2 | ferric |
| pacl-.9 | 4 | 0.2 | aluminum |
| pfc-2.1 | 4 | 0.2 | ferric |
| wac-hb | 4 | 0.2 | aluminum |
| pfc-0.9 | 3 | 0.2 | ferric |
| pfc-1.5 | 2 | 0.1 | ferric |
| kf-4 | 1 | 0.1 | misc |

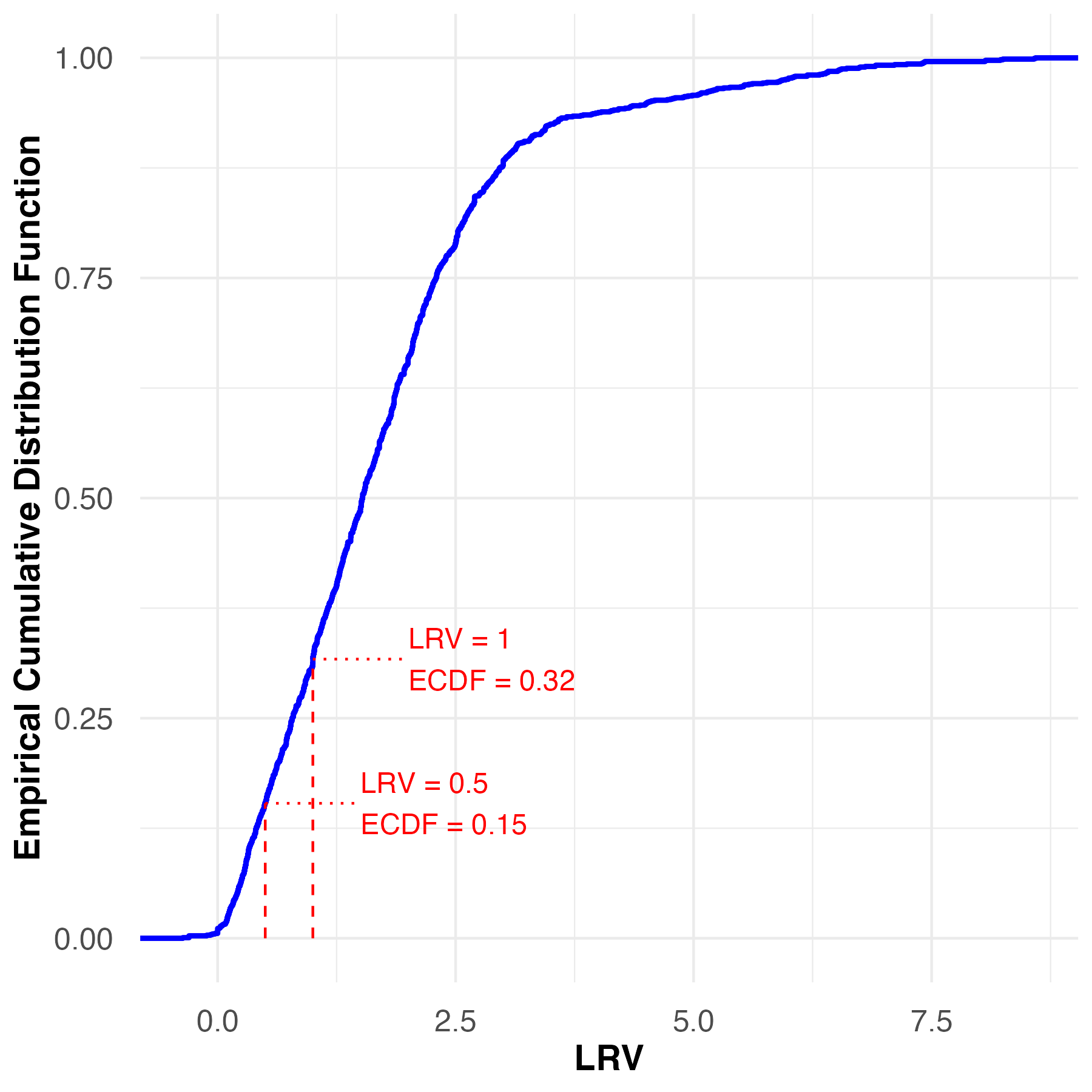

#### **Figure S5:** Empirical cumulative distribution function (ECDF) of all non-censored LRVs from the systematic review

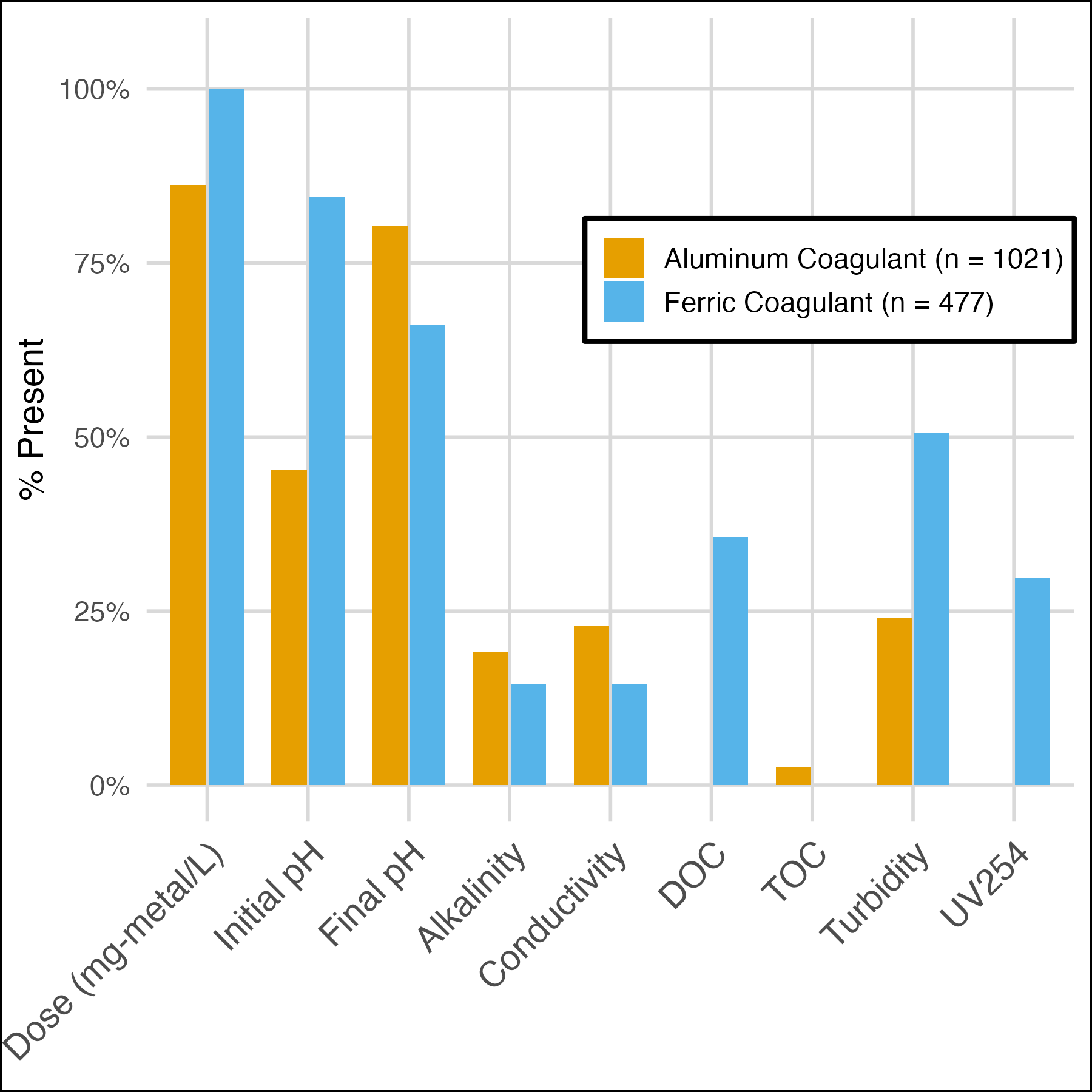

#### **Figure S6:** Percentage of water quality variable values present in the complete data collection for aluminum and ferric coagulants. Includes UV_254_ absorbance, TOC, DOC, conductivity, turbidity, alkalinity, adjusted coagulant dose, and pH (initial and final).

### S9. Model details

#### **Table S14.** Summary of final explanatory model^28^

|  | **LRV.Combined** | | |
| --- | --- | --- | --- |
| *Predictors* | *Estimates* | *CI* | *p* |
| (Intercept) | 2.12 | 1.47 – 2.76 | **<0.001** |
| Treatment Scale [full] | 1.42 | -0.09 – 2.92 | 0.065 |
| Treatment Scale [pilot] | 2.58 | 2.03 – 3.14 | **<0.001** |
| Water Type [secondary effluent] | -1.65 | -2.08 – -1.23 | **<0.001** |
| Water Type [synthetic] | -0.91 | -2.19 – 0.38 | 0.167 |
| Coagulant [aluminum(iii) chloride hexahydrate] | 0.15 | -0.79 – 1.08 | 0.759 |
| Coagulant [ferric chloride] | -0.92 | -1.68 – -0.15 | **0.019** |
| Coagulant [ferric sulfate] | -0.87 | -2.04 – 0.29 | 0.141 |
| Coagulant [ferrous sulfate] | -0.47 | -2.40 – 1.46 | 0.633 |
| Coagulant [pacl] | 1.00 | -0.08 – 2.08 | 0.070 |
| Coagulant [pacl-.9] | -1.05 | -2.34 – 0.23 | 0.107 |
| Coagulantpacl-1.5 | 0.71 | -0.41 – 1.84 | 0.215 |
| Coagulantpacl-1.5ns | 1.10 | -0.04 – 2.23 | 0.058 |
| Coagulantpacl-1.5s | -0.05 | -0.77 – 0.66 | 0.887 |
| Coagulantpacl-1.8s | 1.11 | -0.07 – 2.28 | 0.065 |
| Coagulantpacl-2.1b | 1.83 | 0.90 – 2.76 | **<0.001** |
| Coagulantpacl-2.1c | 2.37 | 1.46 – 3.29 | **<0.001** |
| Coagulantpacl-2.1ns | 1.38 | 0.57 – 2.20 | **0.001** |
| Coagulantpacl-2.1s | 0.75 | -0.06 – 1.56 | 0.068 |
| Coagulantpacl-2.4b | 1.20 | -0.05 – 2.44 | 0.060 |
| Coagulantpacl-2.4c | 1.53 | 0.29 – 2.77 | **0.016** |
| Coagulantpacl-2.7 | 1.65 | 0.57 – 2.73 | **0.003** |
| Coagulantpacl-2.7ns | 2.37 | 1.23 – 3.51 | **<0.001** |
| Coagulantpfc-0.9 | -0.03 | -1.86 – 1.80 | 0.974 |
| Coagulantpfc-1.5 | -1.15 | -3.16 – 0.86 | 0.262 |
| Coagulantpfc-2.1 | -0.90 | -2.82 – 1.01 | 0.355 |
| Coagulant Aid Present | 0.56 | 0.34 – 0.78 | **<0.001** |
| pH after treatment | -0.58 | -0.66 – -0.49 | **<0.001** |
| Virus Measurement Category [Molecular] | -1.31 | -1.47 – -1.15 | **<0.001** |
| log coag dose aluminum | 1.37 | 0.92 – 1.81 | **<0.001** |
| log coag dose ferric | 1.41 | 0.76 – 2.05 | **<0.001** |
| Baltimore Class [dsDNA] | -0.51 | -0.88 – -0.15 | **0.006** |
| Baltimore Class [dsRNA] | -0.75 | -1.72 – 0.22 | 0.128 |
| Baltimore Class [ssDNA] | -0.64 | -1.19 – -0.08 | **0.024** |
| N _Record.ID_ | 20 | | |
| N _Coagulant_ | 22 | | |
| N _Virus.Abbreviation_ | 28 | | |
| Observations | 1050 | | |

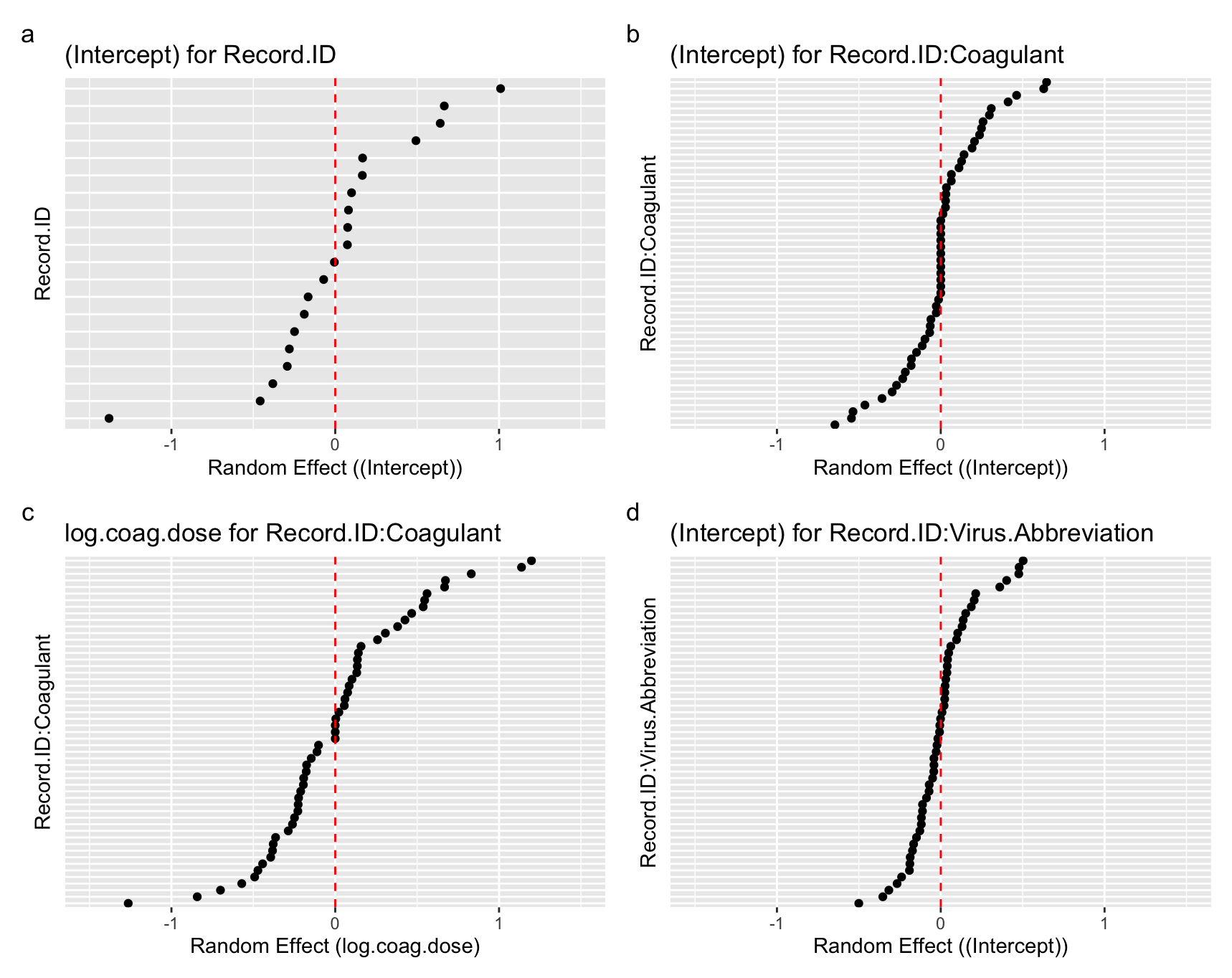

#### **Figure S7.** Random effect structure in the final explanatory model for a) the intercept for paper ID, b) the intercept for coagulant within paper, c) the slope for log_10_(coagulant dose) within paper, and d) the intercept for virus within paper.

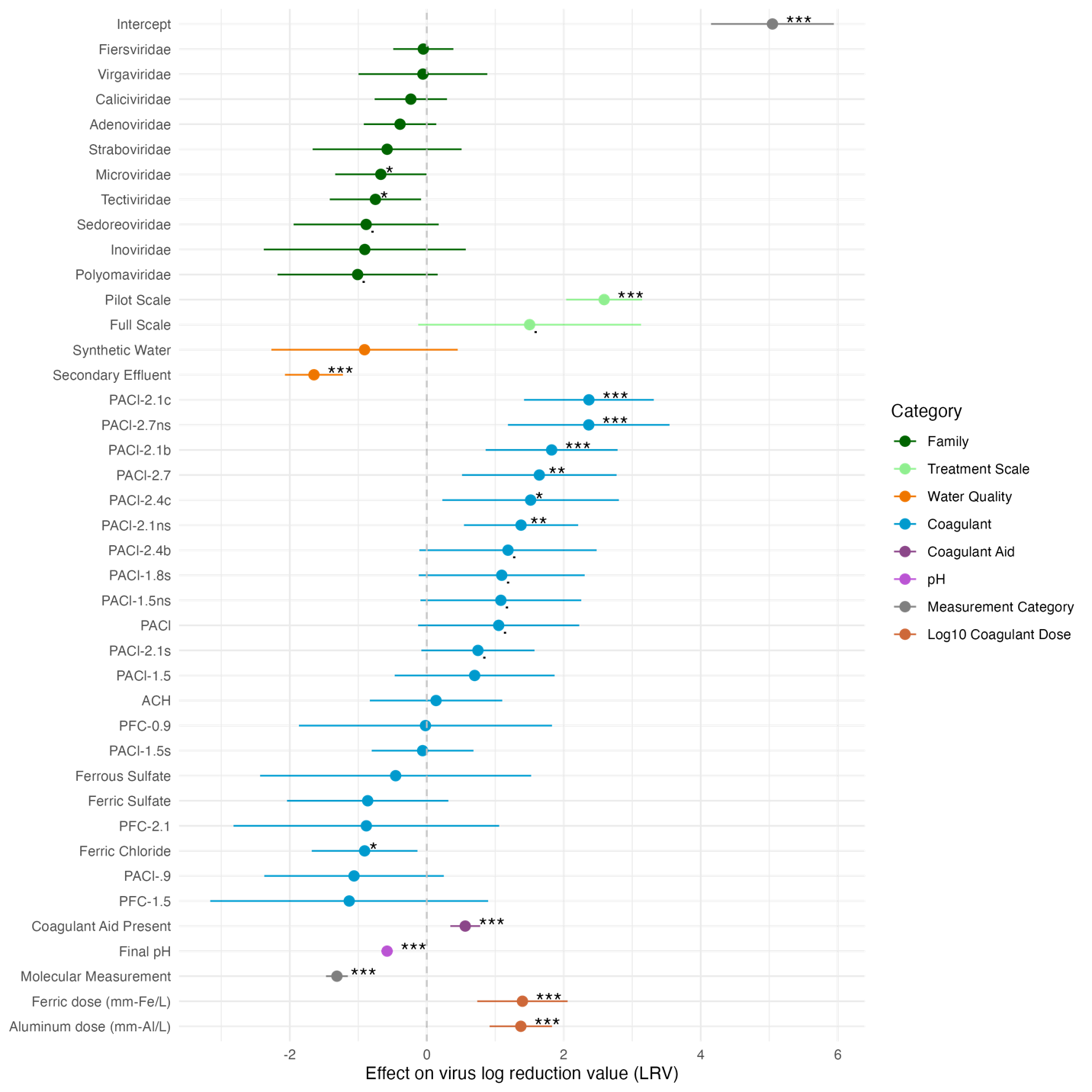

#### **Figure S8.** Coefficients for categorical and numerical variables in the alternate taxonomic model. Asterisks indicate statistical significance (“.”: < 0.1; * : < 0.05; **: < 0.01; ***: < 0.001). Points show the impact of the variable on model LRV estimates relative to a reference value for each variable; the addition of the values of these coefficients to the model intercept provides the estimated LRV. The categorical variable reference categories are: aluminum sulfate for the coagulant type variable, coagulant aid absent for the coagulant aid variable, Picornaviridae for the virus family variable, infectivity for the measurement variable, bench scale for the treatment scale variable, and drinking water source for the water quality variable. For numerical variables, log10(coagulant dose) was referenced to 0 (coagulant dose of 1 mg-metal/L), and pH was referenced to pH 5.

#### **Table S15** Summary of taxonomic model

|  | **LRV.Combined** | | |
| --- | --- | --- | --- |
| *Predictors* | *Estimates* | *CI* | *p* |
| (Intercept) | 5.05 | 4.16 – 5.93 | **<0.001** |
| Family [Adenoviridae] | -0.39 | -0.89 – 0.11 | 0.129 |
| Family [Caliciviridae] | -0.23 | -0.74 – 0.28 | 0.372 |
| Family [Fiersviridae] | -0.05 | -0.47 – 0.37 | 0.816 |
| Family [Inoviridae] | -0.90 | -2.36 – 0.55 | 0.223 |
| Family [Microviridae] | -0.67 | -1.30 – -0.04 | **0.038** |
| Family [Polyomaviridae] | -1.01 | -2.15 – 0.13 | 0.083 |
| Family [Sedoreoviridae] | -0.88 | -1.91 – 0.14 | 0.091 |
| Family [Straboviridae] | -0.58 | -1.60 – 0.44 | 0.264 |
| Family [Tectiviridae] | -0.75 | -1.38 – -0.12 | **0.020** |
| Family [Virgaviridae] | -0.06 | -0.97 – 0.86 | 0.904 |
| Treatment Scale [full] | 1.50 | -0.05 – 3.06 | 0.058 |
| Treatment Scale [pilot] | 2.59 | 2.04 – 3.14 | **<0.001** |
| Water Type [secondary effluent] | -1.65 | -2.07 – -1.22 | **<0.001** |
| Water Type [synthetic] | -0.91 | -2.21 – 0.39 | 0.172 |
| Coagulant [aluminum(iii) chloride hexahydrate] | 0.14 | -0.80 – 1.07 | 0.777 |
| Coagulant [ferric chloride] | -0.91 | -1.67 – -0.14 | **0.020** |
| Coagulant [ferric sulfate] | -0.86 | -2.02 – 0.30 | 0.145 |
| Coagulant [ferrous sulfate] | -0.45 | -2.36 – 1.45 | 0.640 |
| Coagulant [pacl] | 1.05 | -0.07 – 2.17 | 0.066 |
| Coagulant [pacl-.9] | -1.06 | -2.34 – 0.22 | 0.105 |
| Coagulantpacl-1.5 | 0.70 | -0.43 – 1.82 | 0.223 |
| Coagulantpacl-1.5ns | 1.08 | -0.05 – 2.21 | 0.061 |
| Coagulantpacl-1.5s | -0.06 | -0.78 – 0.65 | 0.868 |
| Coagulantpacl-1.8s | 1.09 | -0.08 – 2.27 | 0.067 |
| Coagulantpacl-2.1b | 1.82 | 0.89 – 2.75 | **<0.001** |
| Coagulantpacl-2.1c | 2.37 | 1.45 – 3.28 | **<0.001** |
| Coagulantpacl-2.1ns | 1.38 | 0.56 – 2.19 | **0.001** |
| Coagulantpacl-2.1s | 0.75 | -0.06 – 1.55 | 0.069 |
| Coagulantpacl-2.4b | 1.19 | -0.06 – 2.43 | 0.062 |
| Coagulantpacl-2.4c | 1.52 | 0.28 – 2.76 | **0.017** |
| Coagulantpacl-2.7 | 1.64 | 0.56 – 2.72 | **0.003** |
| Coagulantpacl-2.7ns | 2.36 | 1.23 – 3.50 | **<0.001** |
| Coagulantpfc-0.9 | -0.02 | -1.84 – 1.81 | 0.984 |
| Coagulantpfc-1.5 | -1.13 | -3.14 – 0.87 | 0.268 |
| Coagulantpfc-2.1 | -0.88 | -2.79 – 1.03 | 0.365 |
| Coagulant Aid Present | 0.56 | 0.34 – 0.78 | **<0.001** |
| pH after treatment | -0.58 | -0.66 – -0.49 | **<0.001** |
| Virus Measurement Category [Molecular] | -1.31 | -1.47 – -1.15 | **<0.001** |
| log coag dose aluminum | 1.37 | 0.93 – 1.82 | **<0.001** |
| log coag dose ferric | 1.40 | 0.76 – 2.04 | **<0.001** |
| N _Record.ID_ | 20 | | |
| N _Coagulant_ | 22 | | |
| N _Virus.Abbreviation_ | 28 | | |
| Observations | 1050 | | |

### **S10.** pH analysis

Isoelectric points (IEPs) used in the analysis were gathered primarily from a review of virus IEPs.^29^ When this study reported ranges or multiple estimates for a virus IEP, we calculated the average and cross-referenced the values reported with additional literature. IEPs used in analysis are reported in Table 2. Additional literature sources were used to refine IEP values for MS2,^30^ Qβ,^31^ and poliovirus Sabin.^32^ No supporting literature was found for the IEP of Coxsackievirus B6 Schmitt, so the minimum and maximum IEPs of 4.8 and 6.75 reported for different coxsackieviruses were averaged.

To isolate the effects of pH on virus reduction, we identified 66 data subsets (538 LRVs) from the modeling data collection. Subsets were defined by varying pH after treatment while holding constant virus type, treatment scale, measurement method, water type, coagulant, coagulant aid, and coagulant aid dose. 26 subsets with significant LRV–dose correlations (p < 0.1) were excluded in IEP analysis, since their negative LRV–pH slopes could not be distinguished from dose effects, as exemplified in **Figure S10**.

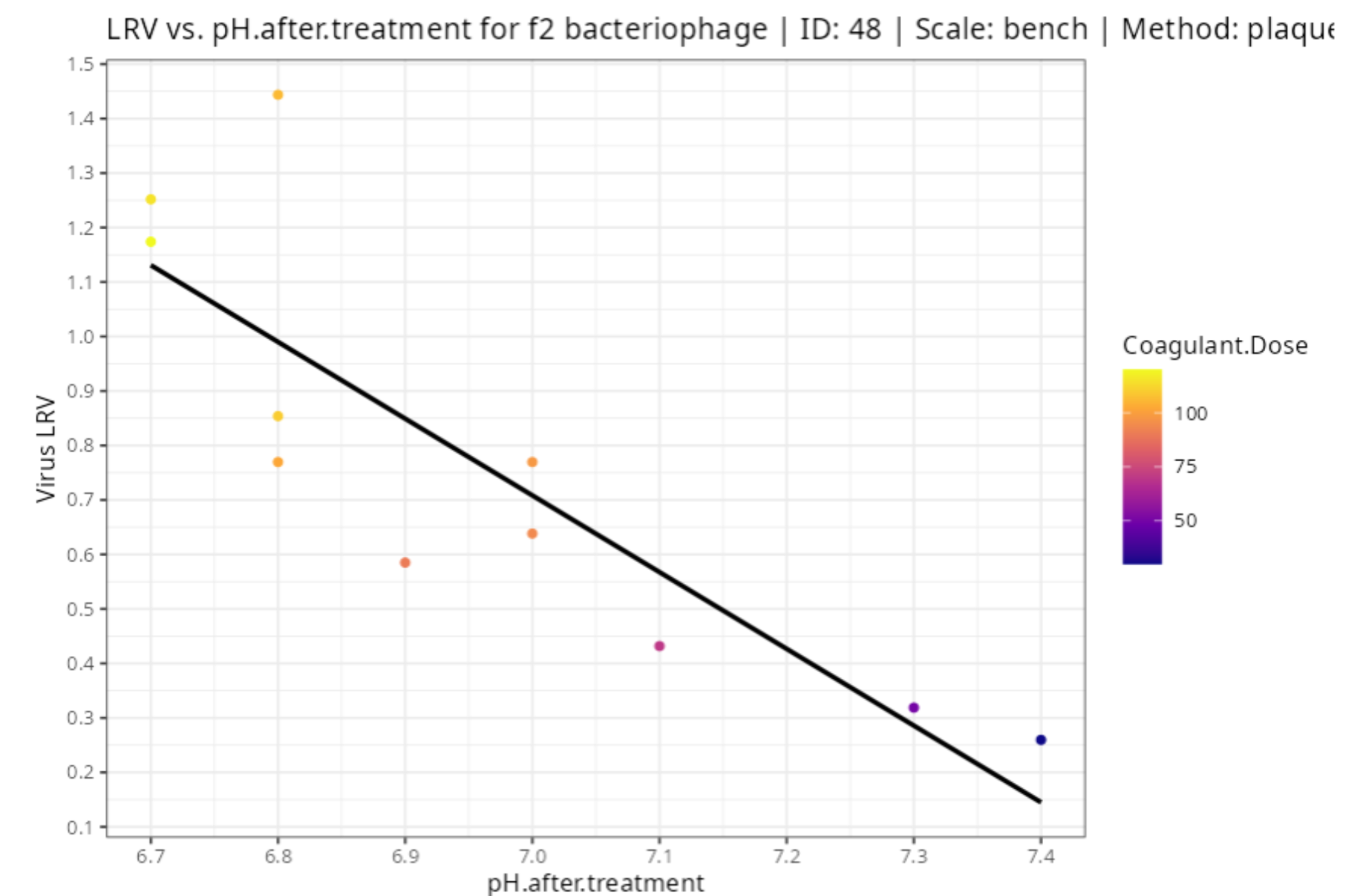

#### **Figure S9.** Example subset showing a strong correlation between coagulant dose and virus reduction. Such apparent correlation with pH led to exclusion of subsets with significant (p < 0.1) LRV-dose relationship.
